# Structure of a sparsely populated chimeric intermediate that facilitates fold-switching of a metamorphic protein

**DOI:** 10.64898/2026.04.23.720511

**Authors:** Bodhisatwa Nandi, Ashok Sekhar

**Author notes:** Corresponding author: Correspondence to: Ashok Sekhar, Molecular Biophysics Unit, Indian Institute of Science Bangalore, Bengaluru-560012, Karnataka, India.

## Abstract

Metamorphic proteins challenge the structure-function paradigm by switching between distinct folds. However, the conformational states sampled along the fold-switching pathway have remained elusive because they are transient and unstable. Using multinuclear chemical exchange saturation transfer and relaxation dispersion nuclear magnetic resonance, combined with chemical shift–based structure determination and protein engineering, we visualized the atomic-resolution structure of an ‘invisible’ state of the metamorphic chemokine lymphotactin. This state is a chimera that integrates the secondary and tertiary structure of one fold with the quaternary assembly of the other. Locking lymphotactin into a single conformation abolishes access to this state, supporting its role as an on-pathway intermediate in metamorphosis. The stabilization of such hybrid intermediates is likely to have facilitated the evolution of fold switching and represent a powerful strategy for protein design.

## Introduction

The single-sequence-structure-function paradigm has served as the cornerstone of structural biology for over five decades and asserts that the amino acid sequence of a protein encodes a single, unique native structure that is tailored to perform its biological function. Metamorphic proteins are striking exceptions to this paradigm and adopt two distinct native structures in the absence of ligands and cofactors^1,2^ with the two folds interconverting between each other and supporting different functions. Metamorphic proteins have been discovered in diverse biological processes: lymphotactin is a secreted chemokine that induces chemotaxis and shows anti-tumour and antimicrobial activity^3,4^; KaiB is a member of cyanobacterial circadian clock^5^; Mad2 is an integral part of the spindle checkpoint complex^6^; and RfaH is a transcription factor that suppresses premature transcription termination^7^. Metamorphic proteins underscore the inherent energetic and structural plasticity of protein folds and illustrate how a single static structure cannot fully capture the complexity of protein function. The structural polymorphism exhibited by metamorphic proteins makes them invaluable for investigating the interplay between protein structure, dynamics, function and evolution. Moreover, their ability to toggle between multiple conformations has garnered considerable interest recently because of their potential application as regulatory elements in synthetic circuits^8–11^.

Lymphotactin (Ltn, also known as XCL1) is a metamorphic chemokine that switches between a monomeric α+β chemokine fold (Ltnαβ)^12^ and a novel dimeric all-β fold (Ltnβ2) ^13^ (Fig. 1A, 1B). When Ltn is secreted by CD8+ T-cells at the site of inflammation or injury, it binds to glycosaminoglycans (GAGs) on extracellular matrix proteins, generating a concentration gradient of immobilized Ltn directed towards the location of the inflammation^14^. Simultaneously, Ltn recognizes the GPCR XCR1 that is expressed on the surface of XCR1+ dendritic cells^15^. The binding of Ltn to its cognate receptor, in conjunction with the concentration gradient of Ltn, forms a navigation system for chemotaxis that guides the migration of dendritic cells to the site of inflammation^14^. While Ltnβ2 is responsible for binding GAGs^13,16^, Ltnαβ interacts with XCR1^17^, making the dynamic equilibrium between both conformations indispensable for chemotaxis. Intriguingly, other chemokines perform both these functions using a single chemokine conformation without requiring this sophisticated fold-switching modality^18^. Clues into the evolutionary benefits conferred by metamorphosis emerge from the non-canonical role played by Ltn in the immune response, where its antimicrobial repertoire is considerably enlarged by its ability to switch between unrelated folds. For example, the novel dimeric Ltnβ2 fold exhibits anti-HIV activity that Ltnαβ does not possess^16,19^. Ltnβ2 also shows bactericidal activity against *Listeria monocytogenes*, *Escherichia coli* and *Salmonella typhimurium*^4^, while both Ltnαβ and Ltnβ2 conformations elicit antifungal response against *Candida*^20^. The metamorphosis of Ltn may therefore constitute a novel weapon in the armoury of organisms in their battle to overcome microbial infection.

**Fig. 1.**
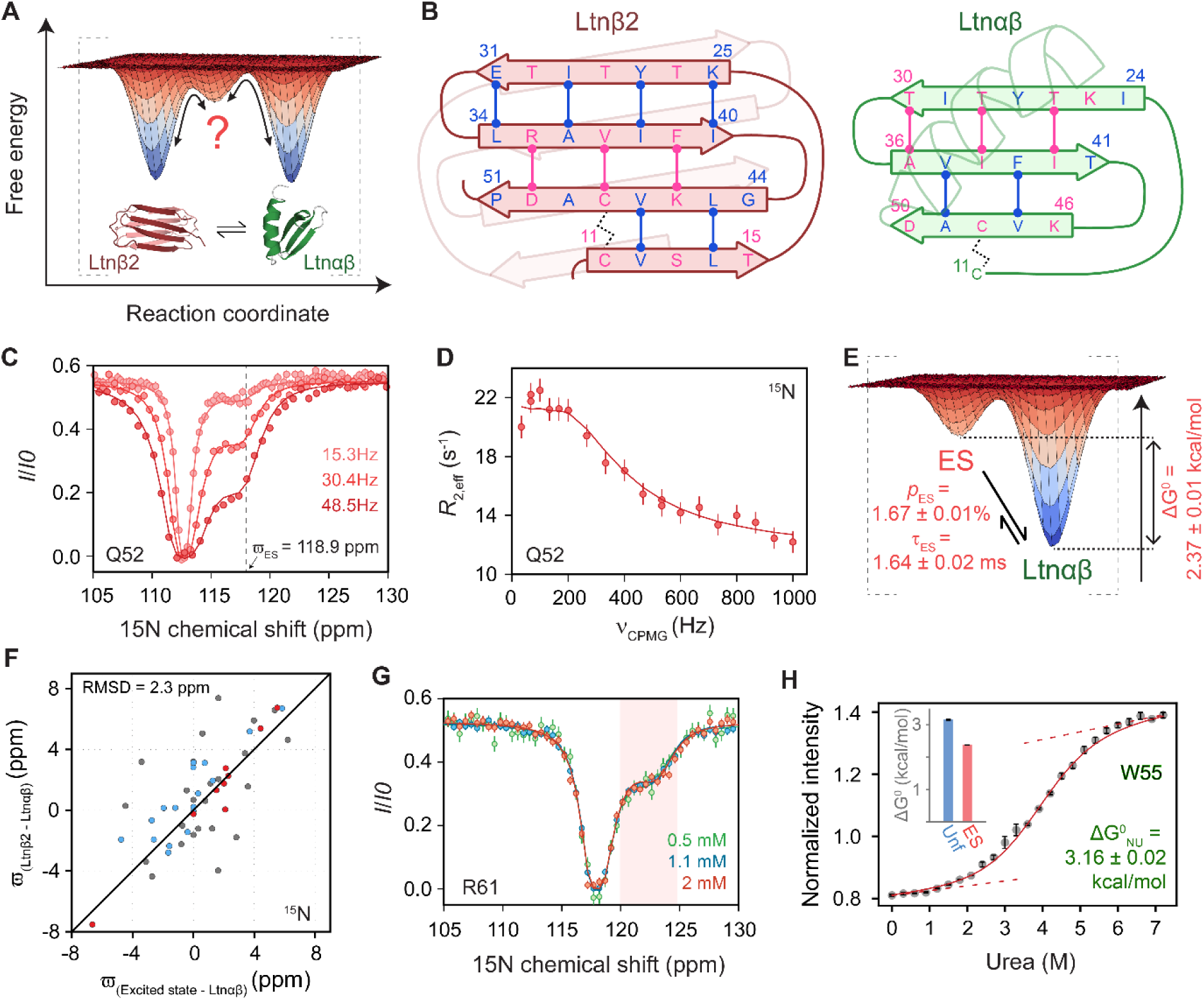
The chemokine conformation of lymphotactin (Ltnαβ) coexists with a partially folded monomeric minor state. **(A)** Schematic of the conformational free energy landscape of Ltn showing a possible on-pathway intermediate for structural interconversion between Ltnαβ (PDB:1J9O)^12^ and Ltnβ2 (PDB:2JP1)^13^. **(B)** Conformational architecture of Ltnαβ and Ltnβ2 displaying hydrogen bonds in the β-sheet. Residues are color-coded depending on whether their sidechains are solvent-exposed (pink) or buried (blue). **(C)** ^15^N CEST profiles of Q52 acquired using multiple radiofrequency field strengths and **(D)** the ^15^N CPMG profile of Q52. Solid lines in both panels represent the global fit of the CEST and CPMG data to the two-state Bloch-McConnell equations and the dashed line in the CEST profile denotes the chemical shift position of Q52 in the excited state. **(E)** Schematic of the conformational free energy landscape showing the free energy difference between Ltnαβ and the ES. The relative population and lifetime of ES are indicated on the plot. **(F)** Comparison of the ^15^N chemical shift difference between the excited state and Ltnαβ (x-axis) with that between Ltnβ2 and Ltnαβ (y-axis). Data points are color-coded based on the secondary structures of the residues in the Ltnαβ conformation: grey: loops, blue: β-strands and red: α-helix. Ltnβ2 chemical shifts at 20 °C were determined by extrapolating the shifts measured at 37.6, 34.7, 30.5, and 28.6 °C. **(G)** ^15^N CEST profiles of R61 at various total concentrations of Ltn. **(H)** Fluorescence-detected urea denaturation of Ltn in 200 mM NaCl at 20 °C, fit to the equation detailed in supplementary text. Ltnβ2 is not detectable under these solution conditions at the start of the titration. The standard free energy change for unfolding of Ltnαβ is indicated in the plot. (Inset) Bar plot comparing the standard free energy changes for unfolding (Unf, blue) of Ltnαβ and its transition to the ES (orange).

Despite the fundamental importance of metamorphic proteins and their widespread applications in biotechnology and cellular engineering, the mechanism of fold-switching remains poorly understood (Fig. 1A). Analogous to the intermediates that guide the *de novo* folding of an unfolded polypeptide to its folded native conformation, discrete high-energy states are likely to be essential for metamorphic proteins to navigate the complexity associated with the remodelling of one folded structure into another. However, experimentally characterizing these intermediates remains challenging due to their short lifetimes and low populations^21^. Solution NMR spectroscopy has recently emerged as a powerful tool for detecting transiently populated states, determining their structure, characterizing their function and elucidating their role in biochemical reaction pathways^22,23^. The potential of NMR for probing such otherwise ‘invisible’ states has opened up opportunities for augmenting our understanding of protein metamorphosis^24–27^. In this report, we use a synergistic combination of multinuclear chemical exchange saturation transfer (CEST)^28^ and Carr-Purcell-Meiboom-Gill relaxation dispersion (CPMG)^29,30^ NMR spectroscopy to determine the atomic resolution structure of a sparsely populated ‘excited’ state coexisting with the chemokine conformation of Ltn. Structural analysis reveals that the excited state shares features of both the Ltnαβ and Ltnβ2 conformations. Experiments carried out on a kinetically trapped variant of Ltn provide compelling evidence that the excited state is an on-pathway intermediate that mediates metamorphosis in Ltn. The stabilization of this chimeric intermediate may also have been important in the evolution of metamorphosis in Ltn and promises to be a potent design strategy that can be generalized to other proteins for the bottom-up development of conformational switches.

## Results

### Ltnαβ transiently populates a higher free energy conformation

Lymphotactin (Ltn) belongs to the XC family of chemokines that are unique in having only one intrachain disulfide bond^31^. Ltn is a 93-amino acid protein that reversibly interconverts between the monomeric chemokine fold (Ltnαβ), where the backbone traces an α/β structure, and a novel dimeric fold in which each protomer is made up of a four-stranded β sheet (Ltnβ2)^13^ (Fig. 1B). Under physiological conditions (37 °C and 150 mM NaCl), the two folds have comparable stability and interconvert rapidly (t_ex_ ∼ 1 s)^13^. However, the position of this equilibrium is sensitive to variations in temperature and ionic strength^13^. Structurally, while the C-terminus of Ltn is intrinsically disordered in both Ltnαβ and Ltnβ2, the disordered N-terminus of Ltnαβ forms the fourth strand of the β-sheet in Ltnβ2^13^. Although the boundaries for the other three β-strands remain approximately the same in both Ltnαβ and Ltnβ2, there is a single residue shift in registry between the β-sheets of the two structural polymorphs (Fig. 1B)^13^. The hydrophobic core of the chemokine fold contains residues such as Phe39 and Val59, which are solvent-exposed in the dimer, while the residues that are accessible in the chemokine fold, including Val12 and Ile38, become buried in the dimer. This dramatic rearrangement effectively turns the protein inside-out during fold switching. It is noteworthy that this elaborate structural reorganization occurs against the backdrop of an intrachain disulfide bond between Cys11and Cys48, even though disulfide bonds are typically expected to confer rigidity to the conformational landscape^32,33^.

We first targeted the conformational free energy landscape in the vicinity of Ltnαβ by carrying out experiments in 200 mM NaCl at 10-20 °C, where Ltn predominantly exists in the Ltnαβ conformation^34^. We then “searched” for sparsely populated conformations (referred to here as excited or minor states) coexisting with Ltnαβ using chemical exchange saturation transfer (CEST)^28^ and Carr-Purcell-Meiboom-Gill relaxation dispersion (CPMG)^29,30^ NMR approaches. In both the CEST and the CPMG experiment, minor conformations that are invariably ‘invisible’ in NMR spectra, are visualized through modulations in intensity of the major conformation^28–30^. In the ^15^N CEST experiment^35^, a weak radiofrequency field of amplitude B_1_ is applied at a specific frequency within the expected range of ^15^N chemical shifts for a duration T_ex_ and the intensity of each major state peak (I) is recorded through a ^1^H-^15^N correlation spectrum. The CEST profile is the ratio of I over I_0_ (where I_0_ is the intensity of the same peak in a reference experiment that lacks the T_ex_ period) plotted as a function of the chemical shift at which the RF field is applied. In systems displaying conformational dynamics on the millisecond timescale, the CEST profile has two dips in intensity, a large dip at the chemical shift of the major state (major dip), and a small dip at the chemical shift of the minor (excited) state (minor dip). In the ^15^N CPMG experiment^36^, trains of 180° pulses are applied at increasing repetition rates (ν_CPMG_) on transverse magnetization of the nucleus of interest for a constant-time duration T_ex_. The effective transverse relaxation rate constant (*R*_2,*eff*_) during T_ex_ is quantified through a ^1^H-^15^N correlation spectrum and plotted against ν_CPMG_ in the CPMG profile. For systems undergoing micro-millisecond timescale exchange, there is a change in *R*_2,*eff*_with ν_CPMG_ (referred to as a dispersion), while CPMG profiles in the absence of conformational exchange are flat.

^15^N CPMG profiles of lymphotactin in the Ltnαβ conformation acquired at both 10 °C and 20 °C show clear fingerprints of conformational dynamics (Fig. S1). Since the exchange-induced dispersion in *R*_2,*eff*_is larger at 20 °C, dynamics can be quantified with higher precision; thus all subsequent analysis has been carried out at 20 °C. Consistent with CPMG data, ^15^N CEST profiles of several residues distributed across the protein show distinct minor dips and highlight the existence of a minor conformation (ES) coexisting with Ltnαβ (Fig. 1C, 1D and S2). ^15^N CEST data acquired at multiple radiofrequency fields can be globally modelled along with CPMG profiles using a two-state model of the Bloch-McConnell equations^37^ (*χ*^2^ = 1.2), indicating that both experiments are reporting on the same dynamical transition. The energy function for the fit is found to be very sensitive to changes in global parameters and reflects that the parameters of the ES can be obtained reliably from the modelling of CEST and CPMG data (Fig. S3). The population (*p_E_*) of the ES at 20 °C is 1.67 ± 0.01 % and its free energy is therefore 2.37 ± 0.01 kcal/mol higher than the free energy of the Ltnαβ (Fig. 1E), implying that fewer than 2 molecules per 100 of Ltn are present in the ES conformation. The lifetime (τ_E_) of the ES is 1.6 ± 0.02 ms (Fig. 1E), emphasizing the transient nature of the minor conformation. Because of its small population and short lifetime, this ES is ‘invisible’ in the NMR spectrum of Ltnαβ (Fig. S4) and thus cannot be characterized at atomic resolution through traditional biophysical approaches.

### The ES is monomeric and structured

As the first step in structurally characterizing the ES, we explored whether the ES was merely Ltnβ2 whose population has reduced significantly because of the high salt and lower temperature conditions^13,34^. Since chemical shifts are excellent reporters of protein structure, we compared the ^15^N chemical shift differences between the ES and Ltnαβ with the difference that is expected for a transition between Ltnβ2 and Ltnαβ (Fig. 1F). The correlation between the two is very poor (RMSD = 2.3 ppm), indicating that the ES is not Ltnαβ.

The members of the chemokine family are known to oligomerize in a variety of ways^18,38^. Although Ltn does not form dimers via the chemokine fold^34^, we next evaluated the possibility that Ltn retains vestiges of the oligomerization propensity present in its family members. Since oligomerization is not a unimolecular process, the equilibrium composition is dependent on the total protein concentration. Simulated CEST profiles reporting on a dimeric minor state clearly show a steep dependence of the size of the minor dip on protein concentration, indicating that CEST data can be used to detect Ltn oligomerization (Fig. S5). However, ^15^N CEST profiles acquired on samples containing 0.5 mM, 1.1 mM and 2.0 mM Ltn overlay perfectly on each other, demonstrating that the ES is not a dimer or a higher-order oligomer of Ltn (Fig. 1G).

The interconversion between Ltnαβ and Ltnβ2 involves a global rearrangement of the hydrogen bonding pattern, as there is a single residue shift between the registry of the β-strands present in Ltnαβ and Ltnβ2^13^. There is evidence in literature suggesting that Ltn has to unfold for this rearrangement to take place^39^. This raises the question whether the ES is the globally unfolded state of Ltn. In order to determine the stability of the unfolded state under our experimental conditions, we carried out urea denaturation experiments detected through the fluorescence signal of the lone Trp W55 present in Ltn. Urea denaturation profiles of Ltnαβ can be fit well to a two-state transition, and the free energy change for unfolding is 3.16 ± 0.02 kcal/mol (Fig. 1H, S6). This is much larger than the free energy change for the transition between Ltnαβ and the ES (ΔG^0^ = 2.37 ± 0.01 kcal/mol) and indicates that the ES is at least three-fold more populated than the unfolded state (Fig. 1H) in the absence of urea. Taken together, NMR and fluorescence data confirm that Ltn samples a structured intermediate in its free energy landscape in addition to Ltnαβ and Ltnβ2. This result is particularly striking as the presence of the intramolecular disulfide bond between Cys11 and Cys48 would confer additional rigidity to the structural scaffold of Ltn.

### Synchronous disorder-to-order and order-to-disorder switches operate during the transition of Ltnαβ to the ES

Having established that the ES is not globally unfolded, we next sought to characterize the ES in greater structural detail. Backbone chemical shifts are excellent indicators of protein secondary structure^40^. Since the ES is ‘invisible’ in the NMR spectrum, we measured these shifts using CEST and CPMG experiments targeting ^13^Cα^41–43^, ^13^C’^44,45^, ^1^HN^46,47^ and ^1^Hα^48,49^ nuclei. Minor dips in CEST profiles and dispersions in CPMG profiles could be clearly identified for several residues (Figs. 2A-2F, S7-S9). Moreover, all CEST and CPMG data could be fit using the ES populations (*p_E_*) and exchange rate constants (*k_ex_*) measured from ^15^N CEST and CPMG profiles, confirming that all multinuclear dynamics experiments consistently report on the same ES. Overall, 75 ^15^N, 53 ^13^Cα, 62 ^13^C’, 42 ^1^HN and 8 ^1^Hα chemical shifts were obtained for the ES coexisting with Ltnαβ (Table S1).

**Fig. 2.**
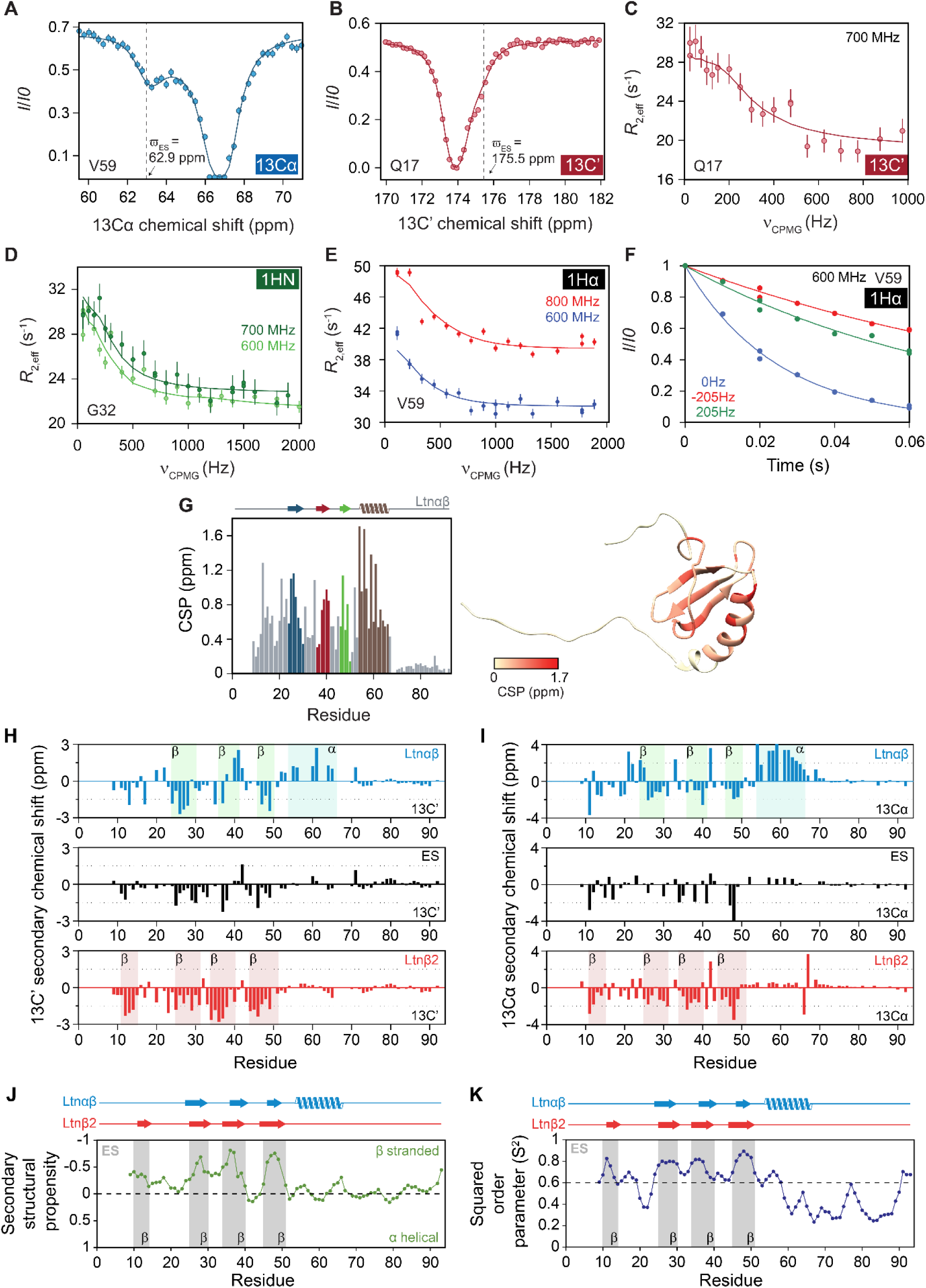
Multinuclear chemical shift information reveals pervasive structural changes across the entire structure of Ltnαβ. CEST profile of the ^13^Cα nucleus of V59 **(A)** and the ^13^C’ nucleus of Q17 **(B)**. **(C)** CPMG profiles of the ^13^C’ nucleus of Q17 acquired at 700 MHz and **(D)** the ^1^HN of G32 acquired at 600 MHz (light green) and 700 MHz (dark green). Solid lines are fits of the data to the two-state Bloch-McConnell equations, and the dashed line in CEST profiles indicates the chemical shift position in the excited state. **(E)** ^1^Hα CPMG profiles of V59 acquired at 600 MHz (blue) and 800 MHz (red). **(F)** ^1^Hα R_1ρ_ profiles of V59 for determining the sign of the chemical shift difference ((|δ_G_|, ν_1_) = (205 Hz, 105 Hz)). **(G)** Residue-specific chemical shift perturbations (CSPs) between Ltnαβ and the ES (left), calculated as described in Methods section. CSPs are depicted as a bar plot (left, color coded according to the secondary structural element present in each region of Ltnαβ) as well as on the structure of Ltnαβ (right, PDB:1J9O)^12^. **(H)** ^13^C’ and **(I)** ^13^Cα secondary chemical shifts of Ltnαβ (top), the excited state (middle) and Ltnβ2 (bottom). The locations of the β-strands and the α-helix in Ltnαβ are shown as light-green and light-blue boxes, respectively, for Ltnαβ, while the β-strands boundaries in Ltnβ2 are shown in light-red boxes. The secondary chemical shifts for Ltnβ2 were calculated using the chemical shifts available in BMRB (ID: 15215). Residue-specific predicted secondary structures **(J)** obtained using the SSP algorithm^54^ and squared order parameters **(K)** derived from backbone chemical shifts using TALOS+^69^. The secondary structure boundaries for Ltnαβ and Ltnβ2 are shown on top of the plots. β-strand boundaries obtained from the CS-Rosetta model of the ES are shown as grey boxes within the plots.

Since chemical shifts are exquisitely sensitive to the local environment of a nucleus, changes in protein secondary and tertiary structure will cause concomitant changes in NMR chemical shifts^50^. In order to determine the regions undergoing the largest change in conformation when Ltnαβ converts to the ES, we first calculated the residue-specific chemical shift perturbation value (CSP), which integrates information from multiple nuclei for each residue into a single score^51^ that correlates with the extent of structural alteration occurring at a site. The CSP plot (Fig. 2G) reveals substantial perturbations in structure occurring across the entire ordered region of Ltn (residues 24–66), while the C-terminus retains its disordered nature in the ES (residues 67–93). Interestingly, although the loops connecting the β-strands show smaller CSPs than the helical and strand regions, the disordered N-terminal region (10–18) preceding β1 exhibits surprisingly large CSPs. This indicates that the structure and dynamics of the N-terminus of Ltnαβ changes during the transition to the ES.

Given that CSPs can be large because of changes in backbone secondary structure or changes in flexibility, we independently assessed the secondary structure of the ES by calculating secondary chemical shifts (SCS)^52,53^, which are deviations of backbone chemical shifts from their random coil values and report on site-specific secondary structure. ^13^C’(Fig. 2H) and ^13^Cα SCS (Fig. 2I) not only confirm that the C-terminus is disordered in the ES but also validate that the helix present between residues 54 and 66 in Ltnαβ has also unfolded during the conversion to the ES, resulting in the large CSPs observed in this region (Fig. 2G). Intriguingly, negative SCS values are seen for ES in segments corresponding to the four β-strands of Ltnβ2 (Fig. 2H, 2I), strongly suggesting that the ES adopts an all-β secondary structure similar to that of Ltnβ2. To provide a more integrated assessment that accounts for the amino-acid-specific variations in secondary chemical shift magnitudes, we calculated secondary structure propensity (SSP)^54^, which combines multiple chemical shifts into a residue-specific score. This score (Fig. 2J) reinforces our previous conclusions and indicates that the N-terminal region between residues 10–15, which is disordered in Ltnαβ, adopts a β-strand conformation in the ES, while the C-terminal helix has unfolded in the ES. In addition, the regions exhibiting high β-strand propensity align better with the strand boundaries of Ltnβ2 rather than Ltnαβ (Fig. 2J) implying that the strand topology of the ES is consistent with the β-sheet of Ltnβ2.

The presence of ordered secondary structure in proteins is expected to reflect in a lack of flexibility. In order to estimate the amplitude of dynamics in the ES, we used the backbone chemical shifts to calculate squared-order parameters^55^ (Fig. 2K), which are less than ∼0.6 in disordered regions and close to 1 in rigid parts of the protein. All the residues showing β-strand propensity in the ES (Fig. 2J) are structurally rigid (S^2^ ∼0.6-0.8); notably, these rigid segments match the strand boundaries of Ltnβ2 and not Ltnαβ, consistent with predictions from SCS and SSP scores. Similarly, residues downstream of Asp58 at the C-terminus have S^2^ values lower than 0.6, demonstrating their intrinsic flexibility and agreeing with the lack of secondary structure in this region. In summary, backbone chemical shifts of the ES measured using CEST and CPMG experiments reveal that the fluctuation of Ltnαβ to the ES involves an order-to-disorder transition of the C-terminal α-helix, and a simultaneous ordering of the disordered N-terminal segment between 10 and 15 into a β-strand, culminating in a four-stranded β-sheet ES.

### The ES is a monomeric four-stranded β-sheet resembling the protomer of Ltnβ2

Although backbone chemical shifts can be analyzed using empirical algorithms to obtain patterns in secondary structure, they do not explicitly define the topological arrangement of these strands. In order to obtain a three-dimensional model for the ES, we next used the CS-Rosetta algorithm^56,57^, which takes backbone chemical shifts as restraints within the Rosetta force^58^ field to model protein structures. The CS-Rosetta calculation of the ES structure using CEST- and CPMG-derived backbone chemical shifts converges robustly as seen from the energy vs RMSD plot (Fig. S10). The ensemble of 10 lowest-energy structures is shown in Figure 3A and a representative model from this ensemble is shown in Figure 3B. The chemical shifts of the ES predicted^59^ from the structural models match well with input chemical shifts (Fig. S11), confirming that the ES structure is a faithful representation of the input experimental data.

**Fig. 3.**
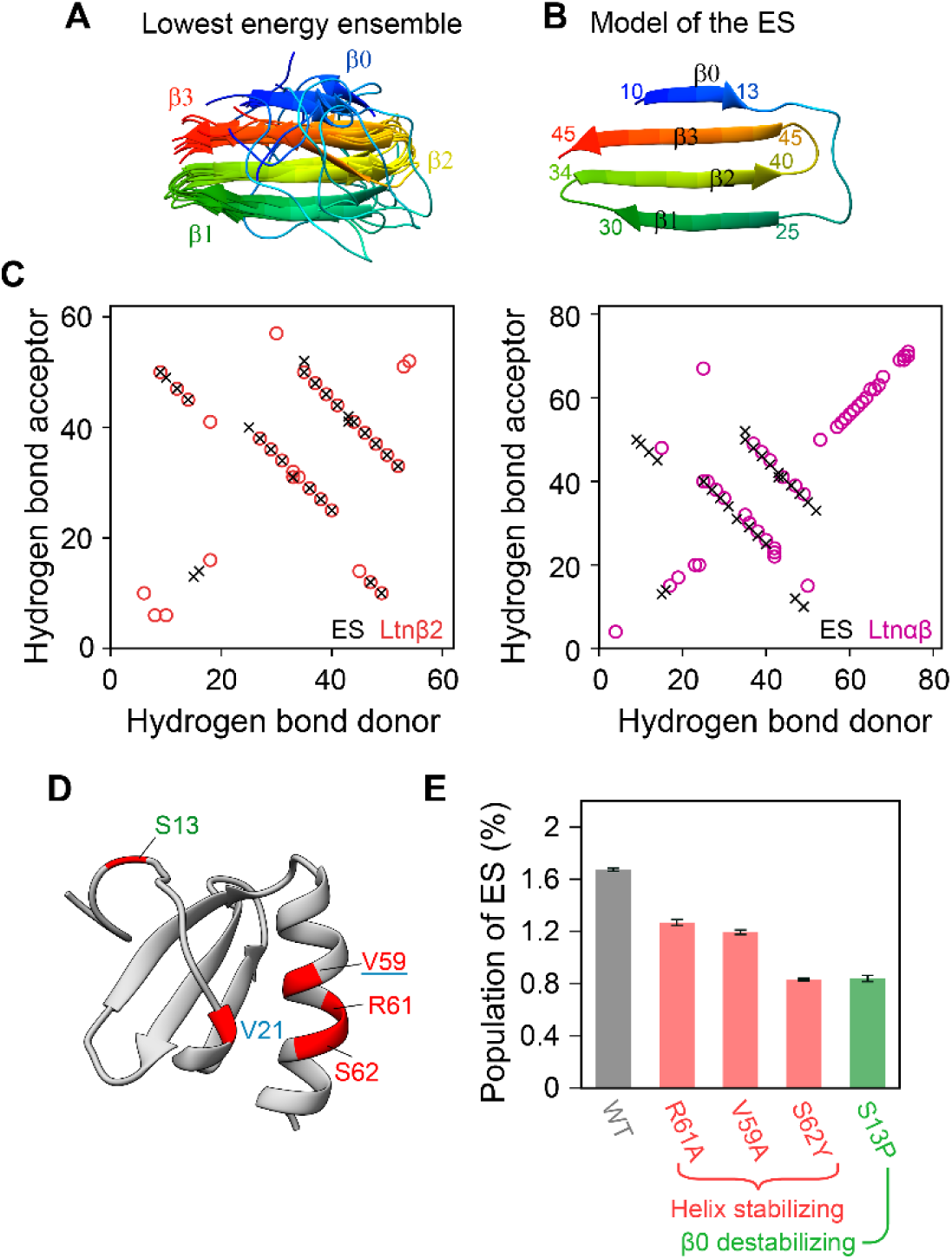
The structure of the excited state. **(A, B)** The structure of the ES calculated using CS-Rosetta. The ensemble of 10 lowest-energy structures **(A)** and a representative model from the ensemble **(B)** are shown in cartoon representation. **(C)** The hydrogen bonding pattern in the ES overlaid with the pattern in Ltnβ2 (left) and Ltnαβ (right). Each plot is a correlation between the residue numbers of the hydrogen bond donor (x-axis) and the hydrogen bond acceptor (y-axis). **(D)** Structure-guided variants designed to stabilize or destabilize the ES, plotted on the structure of Ltnαβ (PDB:1J9O)^12^. V59, R61, and S62 were mutated to stabilize the helix, while S13 was mutated to verify the presence of β0 in the ES. The CC3 variant (V21C/V59C Ltn) was used to probe the exchange mechanism between Ltnαβ and Ltnβ2 and is locked in the Ltnαβ conformation. **(E)** Populations of the ES in variants of Ltn obtained using CEST and CPMG data.

The ES bears a striking resemblance to the protomer of Ltnβ2 dimer and is composed of four β-strands bounded by the residues 10–13, 25–30, 34–40 and 45–51. These β-strands are arranged in the form of an antiparallel β-sheet that overlays much better with the β-sheet of Ltnβ2 than the three-stranded β-sheet in Ltnαβ (Fig. S12). The hydrogen bond network within the ES also matches the pattern in Ltnβ2 and shows the same single residue registry shift that is observed between the hydrogen bond networks of Ltnβ2 and Ltnαβ (Fig. 3C). This structural similarity is further reflected in the observation that the ¹³Cα chemical shifts of the ES agree much more closely with those of Ltnβ2 than with those of Ltnαβ (Fig. S13). A critical consequence of this three-dimensional topology is that the hydrophobic residues present at the core of the Ltnβ2 dimer interface remain exposed to the solvent in the monomeric ES (Fig. S14) and are partly responsible for its lower thermodynamic stability.

We next used an array of complementary strategies to validate the structural features seen in the ES. Notably, the β-strand boundaries defined in the CS-Rosetta structure match well with the rigid regions of ES (Fig. 2K) showing high β-strand propensity (Fig. 2J). In addition to this data-driven validation, we also used structure-guided mutagenesis to systematically stabilize or destabilize specific structural elements observed in the ES (Fig. 3D). First, truncation of the C-terminus (residues 69–93) does not significantly alter the population of the ES (Fig. S15), confirming that the tail neither acquires ordered structure nor contributes to the tertiary fold of the ES. Next, since the C-terminal α-helix (residues 54–66) of Ltnαβ is unfolded in the ES, we designed several mutations such as V59A, R61A and S62Y to stabilize this helix (Fig. 3D). While Ala has an intrinsically high helical propensity^60^, the polar sidechain of Ser62 can compete with backbone carbonyls for hydrogen bonding, and the replacement of Ser by the bulky and non-polar Tyr residue is expected to reinforce the canonical hydrogen bonding pattern of the α-helix. CEST and CPMG datasets indicate that the population of the ES in these helix-stabilizing mutants is lower than that of WT Ltn (Fig. 3E, S16-S18), establishing that the helix unfolds in the ES. Thereafter, we sought to investigate the β-sheet architecture of the ES. Although three of the four β-strands of the ES are present in all CS-Rosetta models, the β-strand between residues 11–14 (β0) is absent in a fraction of the models (Fig. 3A, S19). In order to probe the existence of this β-strand in the ES, we destabilized β0 by replacing Ser13 with Pro (Fig 3D). The imino nitrogen of Pro cannot serve as a hydrogen bond donor, and the pyrrolidine ring restricts the N-Cα dihedral angle ϕ to ∼ −65°, which is unsuitable for a β-strand backbone conformation^61^. The population of the ES in S13P Ltn is two-fold lower than in WT (Fig. 3E, S20), confirming that β0 is indeed present in the ES and contributes to its overall thermodynamic stability. Finally, we attempted to stabilize the ES by destabilizing the α-helix of Ltnαβ (V56G, V59P), truncating the α-helix (LtnΔ(55–93)) or by disrupting the dimeric interface of a variant locked in the Ltnβ2 conformation^16^ (CC5/V12R). The NMR spectra of these variants (Fig. S21) showed fingerprints of varying degrees of conformational exchange broadening, suggesting that the ES is malleable and is likely to be transiently sampling alternate conformations such as the globally unfolded state.

### The ES facilitates metamorphosis of Ltn

The ES of Ltn is chimeric in nature as it incorporates the secondary and tertiary structural arrangement of Ltnβ2 while retaining the quaternary structure of Ltnαβ. This unique hybrid architecture makes the ES a prime candidate for an on-pathway intermediate driving the metamorphic switch between Ltnαβ and Ltnβ2. Consistent with this role, we demonstrated that the ES is accessible from Ltnαβ on the millisecond timescale (Fig. 1E). However, our efforts to visualize the ES through CEST profiles of Ltnβ2 were unsuccessful. In order to address whether the ES is indeed on the fold-switching pathway, or if it is an unproductive off-pathway intermediate of Ltnαβ, mere manipulation of the ES stability would not suffice, as thermodynamic functions such as free energies depend only on the state of the system and do not contain information about pathways. To resolve this challenge, we needed a strategy that uses a geometric constraint to selectively block the pathways of structural rearrangements required for metamorphosis while leaving the local fluctuations of the native state unaffected. We therefore used a kinetically locked mutant of Ltn, V21C/V59C (referred to as CC3)^17^, in which a second intramolecular disulfide bond rigidifies the chemokine conformation and prevents Ltnαβ from accessing the Ltnβ2 conformation. This disulfide bond is present in many monomorphic chemokines^62,63^ and was lost in Ltn during evolution. Since the ground-state structure of Ltnαβ is not perturbed significantly by this disulfide bond (Fig. 4A, 4B), we postulated that if the ES was an off-pathway intermediate associated with the Ltnαβ region of the free energy landscape, it would remain unaffected (Fig. 4C, bottom). Conversely, if the ES was on-pathway along the transition towards Ltnβ2, it will become inaccessible because of the disulfide clamp in CC3 (Fig. 4C, top). Indeed, CEST and CPMG measurements demonstrate that the ES in CC3 is below the detection limit of these experiments (Fig. 4D). This result provides strong support for the ES as an on-pathway intermediate in the metamorphosis of Ltn.

**Fig. 4.**
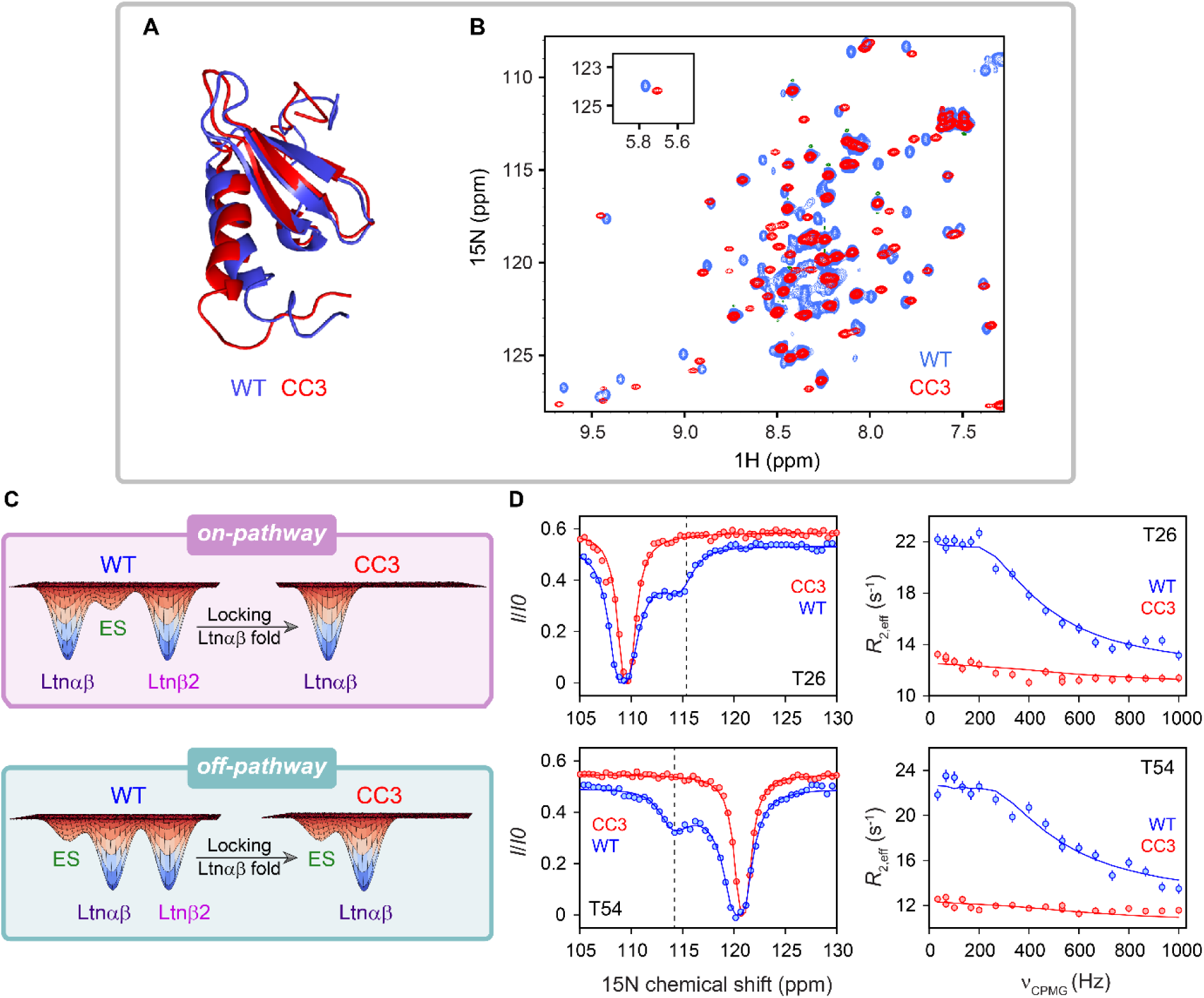
Is the ES on or off the pathway for metamorphosis? **(A)** Overlay of the chemokine structures of WT Ltn (blue, PDB:1J9O)^12^ and the kinetically locked CC3 (V21C/V59C) variant (red, PDB:2HDM)^17^. **(B)** Overlay of the ^15^N-^1^H HSQC spectra of WT Ltn (blue) and the CC3 variant (red). **(C)** Cartoon representation of the strategy for determining whether the ES is on-pathway (top) or off-pathway (bottom). **(D)** Overlay of the CEST and CPMG profiles of T26 (top) and T54 (bottom) in WT Ltn (blue) and the CC3 variant (red). The dashed line in the CEST profiles indicates the chemical shift position of the ES that is observed for WT Ltn.

## Discussion

Metamorphic proteins are referred to as the Janus proteins of structural biology^2^ because of their unique ability to exist in two native structures that can toggle between each other to perform distinct functions. In this report, we use CEST and CPMG NMR experiments to detect an intermediate coexisting with the chemokine fold of Ltn. The intermediate is sparsely populated (p_E_ = 1.67 ± 0.01%) and is short-lived (τ_E_ = 1.64 ± 0.02 ms) and is therefore invisible to traditional biophysical techniques. The atomic resolution structure of this intermediate, visualized through a combination of multinuclear CEST and CPMG data and data-driven structural modelling, reveals that it is chimeric with motifs derived from both Ltnαβ and Ltnβ2. While the secondary structure of the ES and its hydrogen bonding network resemble Ltnβ2, the quaternary structure is monomeric like Ltnαβ. Locking Ltn in the Ltnαβ conformation using the monomorphic CC3 variant also blocks access to the ES. This strongly suggests that the ES must lie on the same pathway connecting Ltnαβ and Ltnβ2. Our results thus highlight that the intermediate facilitating the metamorphosis of Ltn is also Janus-faced, incorporating the structural features of both the conformations favoured by Ltn.

The complexity of the structural rearrangement during the transition between the ES and Ltnαβ suggests that Ltnαβ must first unfold before forming the new set of interactions present in the ES. Indeed, stopped-flow fluorescence spectroscopy studies by Volkman and coworkers concluded that the fold-switching of Ltn must involve global unfolding^39^. Since the population of the unfolded state from urea-denaturation experiments at 20 °C (Fig. 1H) is 0.44 ± 0.02 %, the unfolded state is likely to be invisible even in CEST and CPMG profiles. It is noteworthy that we do not see evidence of the unfolded state in our data. Nevertheless, we still evaluated the CEST and CPMG data using a reaction scheme that is consistent with the findings of Volkman and coworkers and includes an unfolded state between Ltnαβ and the ES (see supplementary text). Interestingly, the population of the unfolded state obtained from this three-state modelling agrees well with the value obtained from our urea titration experiment and the chemical shifts of the ES are consistent with the values determined from the two-state analysis (Fig. S22). Taken together, the model that emerges for the metamorphosis of Ltn is shown in Figure 5A. Ltnβ2 first dissociates to form the ES, which unfolds prior to its reorganization as Ltnαβ. Strikingly, our data reveal that dimerization of the ES is the rate-limiting step for metamorphosis and not fold-switching, which occurs much faster (*k_ex_* = 620 ± 6 s^-1^) than the Ltnαβ↔Ltnβ2 interconversion (∼0.6 s^-1^ at 150 mM NaCl and 25 °C)^64^. This kinetic bottleneck likely arises from the extensive search for a complementary interface that must take place for two ES molecules to bury their exposed hydrophobic surfaces and form the dimer. Given that one of the metamorphic conformations is dimeric in several other metamorphic proteins such as Mad2^6^ and KaiB^5^, our result suggests a shared mechanism of conformational interconversion, where a native state first switches to an unstable monomeric intermediate that exposes hydrophobic patches for intermolecular interactions.

**Fig. 5.**
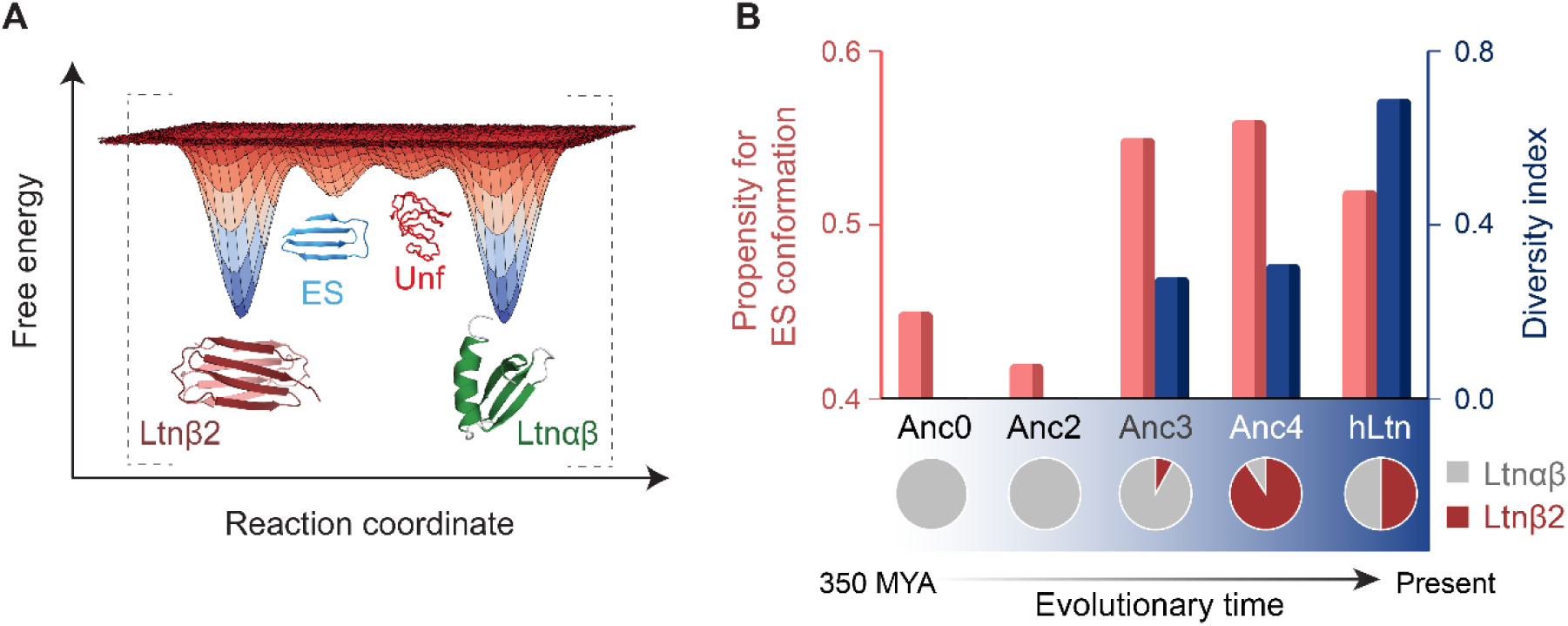
The mechanism of metamorphosis in Ltn. **(A)** Schematic of the proposed free energy landscape of Ltn demonstrating the position of the ES and the probable position of the unfolded state (Unf). **(B)** The propensity of ancestral Ltn sequences^65^ for adopting the ES conformation, estimated using the QMEANDisCO^70^ score as described in supplementary text (red), compared with the diversity index of each ancestral sequence (blue), obtained from the fractional populations of the chemokine (*p*_*C*_) and alternate folds (*p*_*A*_) as the information entropy *S* = −*p*_*C*_ ln *p*_*C*_ − *p*_*A*_ ln *p*_*A*_. The ancestral sequences punctuating the evolutionary trajectory are chronologically plotted on the x-axis and the relative populations of the Ltnαβ and Ltnβ2 conformations for each sequence are shown as a pie chart.

Elegant ancestral reconstruction work^65^ from the Volkman lab has demonstrated that metamorphosis of Ltn was a trait selected for during evolution, and that metamorphic proteins are not mere intermediates in the evolution of protein folds. Dishman et al.^65^ show that the early ancestors of Ltn, Anc.0 and Anc.2, are monomorphic but that Anc.3 (92 % of the chemokine fold) and Anc.4 (9 % of the chemokine fold), which share only 60-70 % sequence identity with Ltn, already have the capacity to switch folds. This suggests that the evolution of the metamorphic traits seen in extant Ltn, where the two folds have comparable thermodynamic stability, may have been preceded by a separate event that was obligatory for the evolution of fold-switching. In order to assess the role of the ES in the evolution of metamorphosis, we threaded the sequences of each of these ancestral proteins in the Ltn lineage onto the structure of the ES. The sequences of both the monomorphic chemokines Anc.0 and Anc.2 are poorly suited for forming the ES (Fig. 5B). Interestingly, however, the stability of the ES in both metamorphic ancestors Anc.3 and Anc.4 is comparable to that of Ltn, despite the alternate fold being present only to 8% in Anc.3. The stabilization of the ES coincides with the initial appearance of metamorphic character in Ltn, when the diversity index of the protein was still low, indicating that fold-switching and dimerization evolved independently in this lineage. It is likely that the ancestral sequences of Ltn first gained the ability to adopt the four-stranded β-sheet structure of ES, and subsequent stabilization of the ES through burial of exposed hydrophobic residues at the dimeric interface eventually resulted in the metamorphic capabilities of extant Ltn. This evolutionary progression aligns with the paradigm proposed by Dan Tawfik^66^, which suggests that new functionalities evolve from pre-existing promiscuous activity. In this context, the transient sampling of the ES structure in ancestral sequences may have provided the promiscuous template necessary for the evolution of the dimeric native state in the Ltn lineage. It is noteworthy that events in the evolutionary trajectory of metamorphosis mirror the sequence in the real-time fold-switching of Ltn, where the transition of Ltnαβ to the ES is followed by the stabilization of the ES as a dimer in the Ltnβ2 fold.

Characterizing the sparsely populated states present in the conformational free energy landscape of metamorphic proteins has remained a challenge because of the inherent heterogeneity in their structural ensembles. Previous studies using CEST and CPMG NMR experiments have been successful in detecting sparsely populated intermediates in metamorphic proteins. Globally and partially unfolded conformations have been shown to coexist with RfaH^26,27,67^ and KaiB^25,68^, while folded minor conformations have been observed in the free energy landscape of Mad2^24^, although a detailed three-dimensional structural model for these excited states are lacking.

In summary, the atomic resolution structure of the Ltn excited state that emerges from our work suggests that chimeric intermediates may represent a unifying feature of protein metamorphosis. The presence of structural features from both conformations enables these chimeras to transition to either fold by using a part of their structure as a scaffold for nucleating the assembly of the rest of the protein. The stabilization of a chimeric intermediate is a potent design principle that is likely operative not only in real-time fold-switching but also in the evolution of metamorphosis in Ltn and could be invaluable in the design and development of conformational switches that are key components in cellular bioengineering.

## Methods

### Expression and purification of unlabeled, U-^15^N, U-^15^N,^13^C and U-^15^N,^13^C/ 50% ^2^H Ltn

The gene encoding human lymphotactin (Ltn) carrying a N-terminal 6x-His tag followed by a linker sequence and TEV-protease site (ENLYFQG) was synthesized by Genscript and cloned into a pET29b(+) plasmid vector. The plasmid was subsequently transformed into *E. coli* BL21(DE3) cells for overexpression. The purification of wild-type (WT) Ltn, as well as the mutants, was performed using a protocol similar to that mentioned in Peterson et al.^71^. Briefly, cells grown either in LB or 2X minimal media^72^ containing ^15^NH_4_Cl (1 g/L of culture) without (U-^15^N) or with (U-^15^N,^13^C) ^13^C glucose (3 g/L of culture) were induced with 1 mM IPTG when the cell culture reached an optical density of ∼ 0.8. For labelling with 50 % ^2^H, cells were grown in minimal media prepared in a mixture of 50% H_2_O and 50% D_2_O containing ^15^NH_4_Cl (1 g/L) and [^13^C_6_,^2^H_7_]-glucose (3 g/L). After 18 hours of induction at 18 °C, cells were harvested and lysed using sonication in lysis buffer (50 mM sodium phosphate, pH 7.4, 300 mM NaCl, 10 mM imidazole, 0.1% (v/v) 2-mercaptoethanol, 1 mM phenylmethylsulfonyl fluoride). The lysate was centrifuged at 13,000 rpm for 40min, and the pellet containing Ltn was redissolved using solubilization buffer (6 M guanidium hydrochloride, 50 mM sodium phosphate, pH 8.0, 300 mM NaCl, 10 mM imidazole, 0.1% (v/v) 2-mercaptoethanol). The resuspension was clarified by centrifugation at 10000 g for 10 min, and the supernatant was passed through a 5 mL Ni-NTA affinity column. The column was washed with the same solubilization buffer having 30 mM imidazole and subsequently, the protein was eluted using 300 mM imidazole. The eluted protein was dialyzed 5 times against 2 L of 0.3% (v/v) acetic acid to remove the denaturant. Formation of the native disulfide bond was accomplished by dropwise addition of the protein solution into 300 mL of oxidation buffer (20 mM Tris, pH 8.0) on a stirrer overnight. Following oxidation, the protein solution was concentrated, and the 6X-His affinity tag was cleaved by the addition of TEV protease in the ratio 1:40 of protein:TEV. Following His-tag cleavage, the native protein was purified using another round of Ni-affinity chromatography followed by a size exclusion chromatography in sodium phosphate buffer (20 mM sodium phosphate, pH 6, 200 mM NaCl, 1 mM EDTA). The purity of the sample was determined by SDS-PAGE, and the identity of the protein was confirmed with mass spectrometry.

### NMR sample preparation

Purified proteins were exchanged into sodium phosphate buffer (20 mM sodium phosphate, 200 mM NaCl, 1 mM EDTA, 0.1% (w/v) sodium azide, pH 6). 10% (v/v) D_2_O were added to the samples for the field-frequency lock. The Ltn sample purified from 50% H_2_O/D_2_O was exchanged into buffer containing 100% D_2_O after purification from the size exclusion column for NMR experiments. The details of the NMR samples used in this work are listed in Table S2.

### NMR data acquisition

All the NMR experiments in this work were carried out on one of the following spectrometers: (1) 14.1 T (600 MHz ^1^H Larmor frequency) Agilent DD2 spectrometer equipped with cryogenically cooled or room temperature single-axis gradient triple resonance probes, (2) 16.5 T (700 MHz ^1^H Larmor frequency) Bruker Avance Neo spectrometer equipped with room temperature TXI or QXI single-axis gradient triple resonance probes and (3) 18.8 T (800 MHz ^1^H Larmor frequency) Bruker Avance III HD spectrometer carrying a room temperature TXI or a cryogenically cooled TCI probe. Post acquisition, spectra were processed in NMRPipe^73^ and visualized using NMRFAM-Sparky software^74^ packages.

#### CEST data acquisition

^15^N^35^, ^13^C’^45^, ^1^H^N46^, ^13^Cα ^41,42^ CEST experiments were acquired on WT Ltn and Ltn mutants using previously published pulse sequences. The details of the experiments are mentioned in Table S3. Weak radiofrequency fields applied during the CEST experiments were calibrated using the method reported in ^75^.

#### CPMG data acquisition

^15^N CPMG experiments were acquired on WT and mutant Ltn samples using the pulse sequence reported in ^36^. ^13^C CPMG datasets were recorded on the carbonyl carbons of WT Ltn using the pulse sequence from literature ^44^. ^1^H^N^ CPMG profiles were acquired on 600 MHz and 700 MHz spectrometers using the pulse sequence reported in ^47^. ^1^Hα CPMG data was acquired on 600 MHz and 800 MHz spectrometers on the WT sample prepared in 50% D_2_O using the pulse sequence reported in ^48^. The experimental details are mentioned in Table S4.

#### R_1_ρ experiment on ^1^Hα

The R_1ρ_ experiments to determine the sign of ^1^Hα chemical shift differences were performed on the 600 MHz spectrometer on a sample of WT Ltn prepared in 50% D_2_O, using the pulse sequence reported in the literature ^49^. The experiment was acquired in a selective 1D manner. The strength and offset of the applied B_1_ field for each residue was estimated by running simulations following the protocol mentioned in ^49^ and are listed in Table S5. For residues with reliable ^1^Hα |δ_G_| values, three R_1ρ_^±,0^ decay curves were obtained with a relaxation delay stretching upto 60 ms.

### Calculating chemical shift perturbations

Residue-specific chemical shift perturbations (CSP) between the ground state and the excited state were calculated by using the expression:

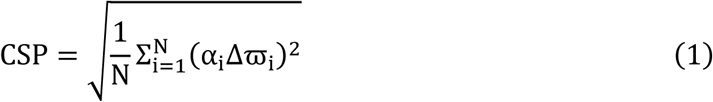

Here, N is the total number of chemical shifts available for a particular residue, α is the scaling factor for each nucleus type and Δϖ is the chemical shift difference between the ground state and the excited state for that particular nucleus. The scaling factor α for a particular nucleus is calculated based on the distribution of its chemical shift in the ground state of Ltn relative to that of the distribution of amide proton chemical shifts^51^. For Ltnαβ, the distributions of ^1^H^N^, ^15^N, ^13^C’, ^13^Cα and ^1^Hα chemical shifts range from 7.45 ppm to 9.72 ppm (2.27 ppm), 108.6 ppm to 127.7 ppm (19.1 ppm), 171.5 ppm to 179.1 ppm (7.6 ppm), 50.4 ppm to 67.3 ppm (16.9 ppm) and 2.7 ppm to 5.8 ppm (3.1 ppm), respectively. Therefore, the scaling factors for ^1^H^N^, ^15^N, ^13^C’, ^13^Cα and ^1^Hα are 1, 2.27/19.1, 2.27/7.6, 2.27/16.9 and 2.27/3.1, respectively. The chemical shift distributions for the ground state are obtained from the BMRB entry of Ltnαβ (accession ID 5042).

### Mutations to perturb ground state and excited state populations

ES structure-guided mutations were designed using a combination of computational tools to selectively stabilize or destabilize either the ES or ground state. To validate the absence of the C-terminal α-helix in the ES, we first aimed to progressively stabilize the helix in the ground state structure, thereby gradually reducing the relative population of the ES. Sequence-based helical propensities were calculated using AGADIR^78^, and changes in the stability of the ground state upon mutations were predicted using FoldX^79^ and PremPS^80^. Firstly, V59A was introduced targeting the central region of the helix, leveraging the known helix-stabilizing effect of the Ala substitution. This strategy was also supported by AGADIR calculations (Fig. S28). The second mutant, R61A, showed increased helicity in AGADIR (Fig. S28) as well as a negative ΔΔG^0^ value in FoldX. The final helix stabilization mutant S62Y, was chosen based on AGADIR (Fig. S28) and yielded negative ΔΔG^0^ values from both FoldX and PremPS. To selectively destabilize the ground state and increase the relative population of the ES, we designed V59P and V56G based on the positive ΔΔG^0^ values predicted by FoldX.

## Acknowledgements

The authors thank Prof. Lewis E. Kay (University of Toronto) for providing the Ltn plasmid and the NMR pulse sequences used in this study and Prof. Pramodh Vallurupalli (TIFR, Hyderabad) for helpful discussions. The authors also thank Claris Niya Varghese and Ahallya Jaladeep for their assistance in running NMR experiments. Support with fluorimeter instrumentation provided by Prof. Mahavir Singh (IISc) is sincerely acknowledged. In addition, the authors thank Dr. Sandhya Sankaran for valuable discussions related to homology modelling. This study made use of NMRbox: National Center for Biomolecular NMR Data Processing and Analysis, a Biomedical Technology Research Resource (BTRR), which is supported by NIH grant P41GM111135 (NIGMS). This work was supported by the DBT/Wellcome Trust India Alliance Fellowship (grant no.: IA/I/18/1/503614), the Ramanujan Fellowship (SB/S2/RJN-167/2017) and a DST/SERB Core Research grant (no. CRG/2019/003457), as well as a start-up grant from IISc awarded to A.S. We also acknowledge funding for infrastructural support from the following programs of the Government of India: DST-FIST, UGC-CAS, and the DBT-IISc partnership program. The Authors would also like to acknowledge the SAIF, DST supported Institute NMR Facility at IISc.

## Author Contribution Statement

Conceptualization: AS

Methodology: BN, AS

Investigation: BN, AS

Visualization: BN, AS

Funding acquisition: AS

Project administration: BN, AS

Supervision: AS

Writing – original draft: BN, AS

Writing – review & editing: BN, AS

