## Supplemental file for "Structure of a sparsely populated chimeric intermediate that facilitates fold-switching of a metamorphic protein"

### Supplementary text

#### Analysis of CEST, CPMG and $R_{1\rho}$ profiles

All the CEST, CPMG (and $R_{1\rho}$) experiments were run in an interleaved pseudo-3D (and pseudo-2D) manner where the change in peak intensities is recorded in the $3^{rd}$ (and $2^{nd}$) dimensions. The peak intensities in CEST and CPMG experiments were subsequently extracted by fitting the lineshape using the program FuDA (https://www.hansen-lab.ucl.ac.uk/fuda/). In some cases, raw intensities were extracted using NMRPipe. For the $R_{1\rho}$ experiment, the intensities were extracted by performing a lineshape analysis in NMRPipe. For the CEST and $R_{1\rho}$ experiments, $I/I_0$ was calculated based on the relative intensity of each plane with respect to a reference plane lacking the exchange duration. In the CPMG experiment, $R_{2,eff}$ values were calculated from $I/I_0$ using the equation $R_{2,eff} = \frac{1}{T_{ex}} \ln \frac{I_0}{I}$ and subsequently plotted against the applied CPMG pulsing frequency.

For WT Ltn, $^{15}$N CEST and CPMG profiles were fit together to the two-state Bloch-McConnell equations[37] using the software package ChemEx (https://gbouvignies.github.io/ChemEx/) to extract fractional populations ($p_E$) and exchange rate constants ($k_{ex}$). CEST profiles acquired at 3 $B_1$ fields (15.3, 30.4, and 48.5 Hz) and CPMG profiles at a $B_0$ field of 700 MHz were globally fit for a total of 14 residues (S13, T15, Q17, R23, K25, T26, F39, R43, V47, Q52, T54, V56, R57, R61) of the WT as these residues have well resolved minor dips in the CEST profiles. The $p_E$ and $k_{ex}$ values obtained in this manner were then fixed while extracting the $^{15}$N excited state chemical shifts for other residues for which either the S/N is low or the minor dip in the CEST profiles is not sufficiently resolved from the major dip. The errors associated with the parameters $p_E$ and $k_{ex}$ were estimated using 1000 Monte Carlo trials (Fig. S3). In the Monte Carlo procedure, synthetic datasets are generated by randomly picking points from a Gaussian distribution, where the mean of the distribution is the best-fit value for every data point and the standard deviation of the distribution is the error in the data points. $\chi^2_{reduced}$ surfaces for $p_E$ or $k_{ex}$ were generated by fitting the datasets while fixing the $p_E$ or $k_{ex}$ to various values.

$^{13}$C\` excited state chemical shifts for WT Ltn were obtained by fitting $^{13}$C\` CEST ($B_1$ = 25Hz) and $^{13}$C\` CPMG profiles ($B_0$ = 700MHz) to the two-state Bloch-McConnell equations while fixing the $p_E$ and $k_{ex}$ to the values obtained from fitting the $^{15}$N CEST and CPMG datasets.

$^{13}$Cα excited state chemical shifts were obtained in a similar manner to $^{13}$C` by fitting 25 Hz $^{13}$Cα CEST profiles acquired on Sample 3 (Table S2) using an out-and-back scheme that transfers magnetization from amide protons to $^{13}$Cα for CEST, before returning back to amide nitrogen for chemical shift labelling and amide proton for detection[41]. $^{13}$Cα CEST data was also acquired on the WT sample prepared in 50% $D_2O$ using a pulse sequence that relies on coherence transfer from $^{1}$Hα to $^{13}$Cα for CEST and back to $^{1}$Hα for detection [42]. For this dataset, the exchange parameters were determined by fitting $^{13}$Cα CEST profiles of V59 acquired at 4 $B_1$ fields (33, 43.8, 54.7 and 65.5 Hz) using a selective-1D scheme [43]. The reliability of the $p_E$ and $k_{ex}$ obtained from fits of V59 was estimated using $\chi^2_{reduced}$ surfaces (Fig. S23). Notably, the $p_E$ and $k_{ex}$ values obtained here differ slightly the numbers obtained using $^{15}$N CEST and CPMG data because the current data was acquired on a partially deuterated sample in 100% $D_2O$ solvent[76]. The $\Delta\varpi$ values for the other residues were then estimated by fitting their 13Cα CEST profiles keeping $p_E$ and $k_{ex}$ fixed to the values obtained from V59.

Most residues in $^{1}H^{N}$ CEST profiles did not exhibit a distinct minor dip, which necessitated the acquisition of $^{1}H^{N}$ CPMG relaxation dispersion experiments at two $B_0$ fields (700 MHz and 600 MHz). The magnitudes of the $^{1}H^{N}$ chemical shift differences($\Delta\varpi_{ES}$) between the ground state (Ltnαβ) and the ES were obtained by fitting $^{1}H^{N}$ CPMG profiles while fixing $p_E$ and $k_{ex}$ to the values determined from the $^{15}$N dataset. Because CPMG profiles do not contain the information on the sign of the chemical shift difference, we sought a systematic method to assign these signs. We observed that the majority of the ES chemical shifts for the backbone nuclei, specifically $^{15}$N, $^{13}$Cα and $^{13}$C' move towards the random coil (RC) values (Fig. S24). Based on this trend, we assumed that the $^{1}H^{N}$ $\Delta\varpi_{ES}$ had the same sign as $\Delta\varpi_{RC}$ ($\varpi_{RC}$-$\varpi_{Ltn\alpha\beta}$). In order to test this assumption, we fit $^{1}H^{N}$ CEST and $^{1}H^{N}$ CPMG data globally for the subset of residues exhibiting a minor dip in their CEST profiles. Both the magnitudes and signs of the $^{1}H^{N}$ chemical shift differences from these fits showed excellent agreement with those derived from the CPMG-only analysis (Fig. S25).

$^{1}$Hα CPMG profiles were fit by fixing the $p_E$ and $k_{ex}$ values obtained by fitting the $^{13}$Cα CEST profile of V59. $^{1}$Hα $R_{1\rho}$ decay profiles were fit to the equation $I = I_0 \, exp(-R_{1\rho}T)$[49]. The signs of $\Delta\varpi$ for $^{1}$Hα excited state chemical shifts were identified based on the faster decaying $R_{1\rho}^{\pm}$ profile.

$^{15}N$ CEST and CPMG profiles of Ltn mutants were fit following protocols similar to those mentioned above for WT Ltn.

**Urea unfolding of Ltnαβ using fluorescence spectroscopy**

Urea titration experiments were performed using three independent sets of samples, each consisting of 21 samples with urea concentrations ranging from 0 to 7.2 M in 0.3 M increments (0,0.3,0.6, …, 7.2 M). The concentration of the urea stock solution was measured using a refractometer, and all samples were prepared in the same buffer as the NMR experiments. The protein concentration in each sample was 9.9 µM for the first two sets and 9.1 µM for the third set. Fluorescence spectra were recorded at 20 °C on an Agilent fluorimeter using an excitation wavelength of 295 nm. Emission was collected from 300 to 500 nm with a scan rate of 52.5 nm/min, averaging time of 0.8 s, and a data interval of 0.7 nm. The emission spectra for all three sets exhibited a red shift at higher urea concentrations. The emission profiles near the maximum were then fit to a 4$^{th}$ order polynomial to precisely determine the value of the emission maximum. The emission maximum for the 0 M urea sample is ~334 nm and for the 7.2 M sample is ~356 nm. Subsequently, the $I_{356nm}/I_{334nm}$ ratio was calculated for every emission profile, and the mean and standard error were determined from the three datasets for each urea concentration. The resulting unfolding profile was then fit to Eq (2) by fixing (Fig. 1H) or fitting the baselines (Fig. S26A) to obtain the free energy of unfolding ($\Delta G^{o}_{NU}$). The reliability of the $\Delta G^{o}_{NU}$ value was then verified by performing a goodness of fit ($\chi^2_{reduced}$) analysis for both fitting protocols (Fig. S6, S26B).

$$I = \frac{\Delta E^{o}_{N} + m_N[D] - \Delta E^{o}_{U} - m_U[D]}{1 + e^{-\frac{\Delta G^{o}_{NU} - m[D]}{RT}}} + \Delta E^{o}_{U} + m_U[D] \quad (2)$$

where I represents the mean ($I_{356nm}/I_{334nm}$), [D] is the urea concentration, and R and T are the gas constant and absolute temperature, respectively. The terms $\Delta E^{o}_{N}$ and $\Delta E^{o}_{U}$ denote the intercepts of the native and the unfolded state baselines, while $m_N$ and $m_U$ indicate their respective slopes. $\Delta G^{o}_{NU}$ is the standard Gibbs free energy of unfolding in $H_2O$ and m characterizes the dependence of $\Delta G^0$ on the buffer.

**Simulating CEST profiles for a homodimerization exchange process**

For two molecules in the ground state (A) conformation, forming a symmetric homodimer ($A_2$) in the excited state, the exchange reaction can be written as:

$$A + A \leftrightharpoons A_2 \tag{3}$$

where [A] and [$A_2$] are their respective concentrations at equilibrium. The dissociation constant $K_d$ can be written as:

$$K_d = \frac{[A]^2}{[A_2]} \tag{4}$$

The total protein concentration $A_T$ is given by the mass balance equation:

$$A_T = 2[A_2] + [A] \tag{5}$$

which can be rewritten as:

$$[A_2] = \frac{A_T - [A]}{2} \tag{6}$$

Combining Eq (4) and Eq (6),

$$K_d = \frac{2 \cdot [A]^2}{A_T - [A]} \tag{7}$$

Rearranging Eq (7),

$$[A] = \frac{-K_d \pm \sqrt{K_d^2 + 4 \cdot 2 \cdot K_d \cdot A_T}}{4} \tag{8}$$

The solution with the negative root holds no physical meaning; hence, taking the positive square root,

$$[A] = \frac{-K_d + \sqrt{K_d^2 + 4 \cdot 2 \cdot K_d \cdot A_T}}{4} \tag{9}$$

The fractional population of the monomeric ground state ($p_A$) and the dimeric excited state ($p_{A_2}$) is given by:

$p_A = \frac{[A]}{A_T}, \quad p_{A_2} = \frac{2 \cdot [A_2]}{A_T}$

Using Eq (4) and Eq (9), $K_d$ was calculated for a $p_{A_2}$ of 1.67 % at a total protein concentration of 1.1 mM to be 127 mM. The $K_d$ value was then used to calculate the $p_{A_2}$ at various total sample concentrations ($A_T$) (Fig. S5A).

For symmetric homodimerization, the exchange rate $k_{ex}$ can be written in terms of the forward ($k_{on}$) and backward ($k_{off}$) rate constants

$$k_{ex} = 2k_{on}[A] + k_{off} \tag{10}$$

$K_d$ is related to $k_{on}$ and $k_{off}$ as:

$$K_d = \frac{k_{off}}{k_{on}} \tag{11}$$

Combining Eqs (10) and (11),

$$k_{on} = \frac{k_{ex}}{2 \cdot [A] + K_d} \tag{12}$$

Using the experimental value of $k_{ex}$ of 620 $s^{-1}$, $k_{on}$ and $k_{off}$ were found using Eqs (7, 8, 11 and 12). Thereafter, these $k_{on}$ and $k_{off}$ values were used to calculate $k_{ex}$ at various sample concentrations and are plotted in Figure S5A.

The $p_{A_2}$ and $k_{ex}$ values at various sample concentrations were then used to simulate the CEST profiles shown in Figure S5B.

**ES structure calculation using CS Rosetta**

Chemical shift-derived squared order ($S^2$) parameter values of the ES (Fig. 2K) indicate that only the segment between residues 10–50 is rigid ($S^2$ ~0.8) and has a well-defined fold in the ES. Consequently, structure calculations were restricted to the region spanning residues 9–52. All structure calculations in this study were performed in NMRbox[77] using CS-Rosetta[56]. The initial calculation was performed using $^{15}N$, $^{13}C\alpha$, $^{13}C'$ and $^{1}H^{N}$ chemical shifts. Firstly, a fragment library was constructed containing 200 3-residue and 200 9-residue fragments for each position within the 9–52 sequence range. During the fragment generation step, Cys11 and Cys48 were explicitly defined to be in the oxidized state by writing the amino acid symbol as lowercase 'c' in the input chemical shift table. This ensured that the selected fragments reflected the local conformational preferences of disulfide-bonded cysteines. To ensure the *de novo* nature of the structure calculation, the resulting fragment library was analyzed for potential structural bias. The library exhibited high statistical diversity, with the most frequent PDB structure accounting for less than 1.6% and 0.4% of the total 3-mer and 9-mer fragments, respectively. Thereafter, fragment assembly was performed using the Rosetta topology broker protocol, generating 5000 structures through the AbinitioRelax application[56,57]. This protocol assembles the chemical shift-derived fragments via a Monte-carlo process followed by an all-atom refinement step. The disulfide bond between Cys11 and Cys48 was held intact during this routine using the detect_disulf and fix_disulf flags. The resulting models were then evaluated using the standard scoring function: $E=E_{Rosetta}+E_{chemical\ shift}/4$ [56]. The 10 structures with the lowest scores were used to define the ES ensemble (Fig. 3A). The Cα RMSD was calculated for all 5000 structures against the lowest-energy structure over the residue range 23–52. A characteristic funnel-shaped energy profile was obtained (Fig. S10) that signifies convergence of the CS-Rosetta structure calculation protocol. The lowest energy ensemble has a Cα RMSD of 2.3 Å for residues 23–52, while the RMSD increases to 6.8 Å when calculated for residues 9–52. This indicates that the N-terminal region has a higher level of conformational flexibility in the structural models of the ES. Addition of $^{1}$Hα chemical shifts to the structure calculation did not significantly alter the final ensemble (Fig. S27A). In order to test whether the structural features of the lowest energy ensemble depend on the residue boundaries or sampling depth, we calculated another ensemble for residues 9–54, generating a total of 50,000 structures (Fig. S27B); the lowest energy structures from this calculation had the same structural motifs seen in the ensemble generated for residues 9–52.

**3-state fitting of CEST and CPMG data to accommodate the unfolded state**

$^{15}$N CEST and CPMG data were fit to a 3-state model that includes the unfolded (Unf) state as an on-pathway intermediate between the Ltnαβ and ES. CEST and CPMG profiles for residues S13, T15, Q17, R23, K25, T26, F39, R43, V47, Q52, T54, V56, R57, and R61 were globally fit to the Ltnαβ↔Unf ↔ES model, with the unfolded state chemical shifts fixed to the values obtained for Ltn sequence using ncIDP[81] (Fig. S22A). The free energy difference between Ltnαβ and the unfolded state derived from this three-state fit ($\Delta G^0$=2.89 ± 0.04 kcal/mol) is in close agreement with the free energy of unfolding ($\Delta G^0_{NU}$=3.16 ± 0.02 kcal/mol) of Ltnαβ determined independently through urea titration experiments (Fig. S22B). Notably, the derived ES chemical shifts are largely invariant to the model (two-state or three-state) used for fitting CEST and CPMG data (Fig. S22C).

**Assessment of Conformational Preferences for the ES Fold in Ancestral Lymphotactin Sequences**

We evaluated the ability of the Ltn ancestral sequences[65] to adopt the ES conformation using homology modelling. Each of the Ancestral sequences (Anc0, Anc2, Anc3 and Anc4), along with the Ltn sequence, was modelled into the ES conformation using SWISS-MODEL[82], using the four-β-stranded ES structure as a template. To quantitatively evaluate the fitness of these sequences for the ES fold, we monitored the QMEANDisCO score[70]. Thereafter, to assess the evolution trajectory of the ES fold in lymphotactin, we compared the scores of the ancestral sequences against that obtained for the Ltn sequence.

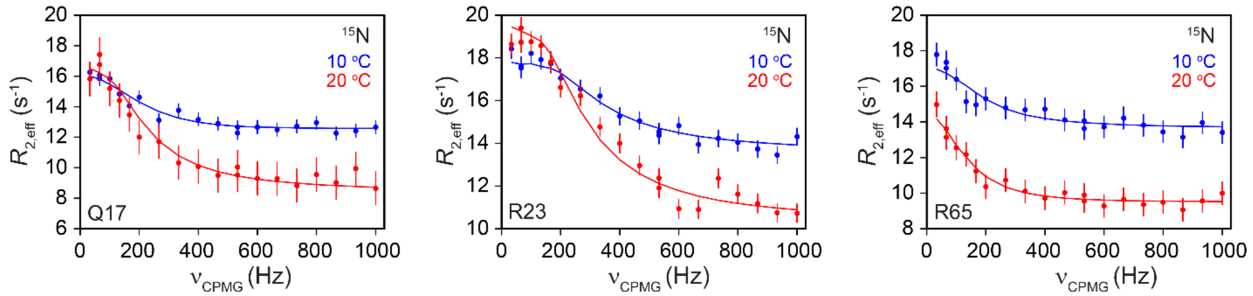


**Fig. S1. Temperature dependence of the Ltnαβ↔ES exchange process.** $^{15}N$ CPMG profiles acquired on Ltn at 10 °C (blue) and 20 °C (red). CPMG data was acquired on a 700 MHz spectrometer.

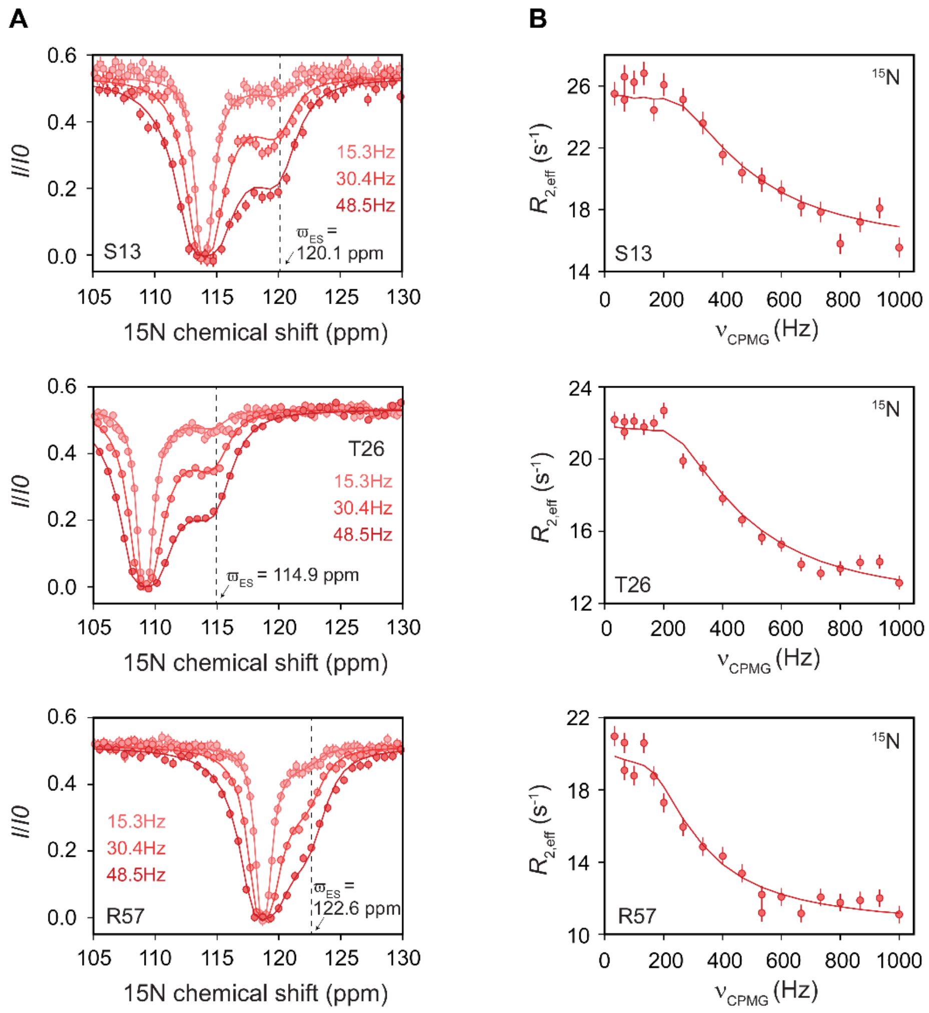


**Fig. S2. The Ltnαβ↔ES exchange process is visible at multiple sites in Ltnαβ.** A) CEST profiles of S13, T26 and R57 recorded at three $B_1$ fields. B) CPMG profiles of S13, T26 and R57 recorded on a 700 MHz NMR spectrometer.

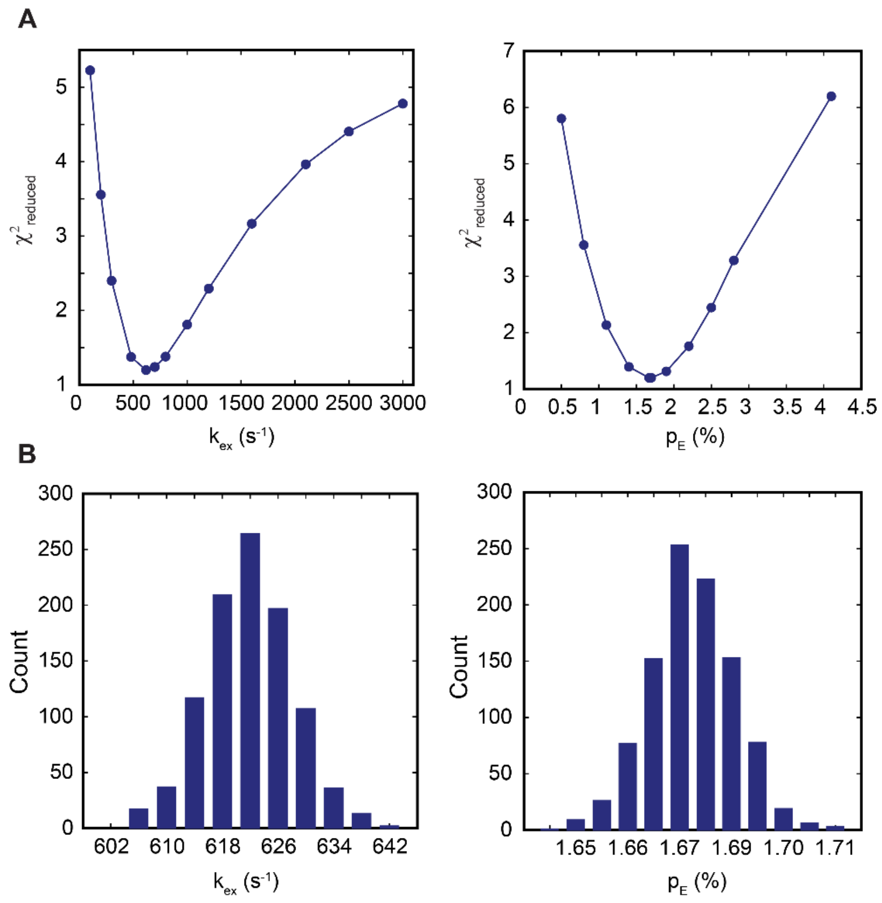


**Fig. S3. Statistical analysis of the exchange parameters for Ltnαβ↔ES exchange.** A) $\chi^2_{reduced}$ surfaces for $k_{ex}$ (left) and $p_E$ (right). B) Monte Carlo parameter distributions for $k_{ex}$ (left) and $p_E$ (right).

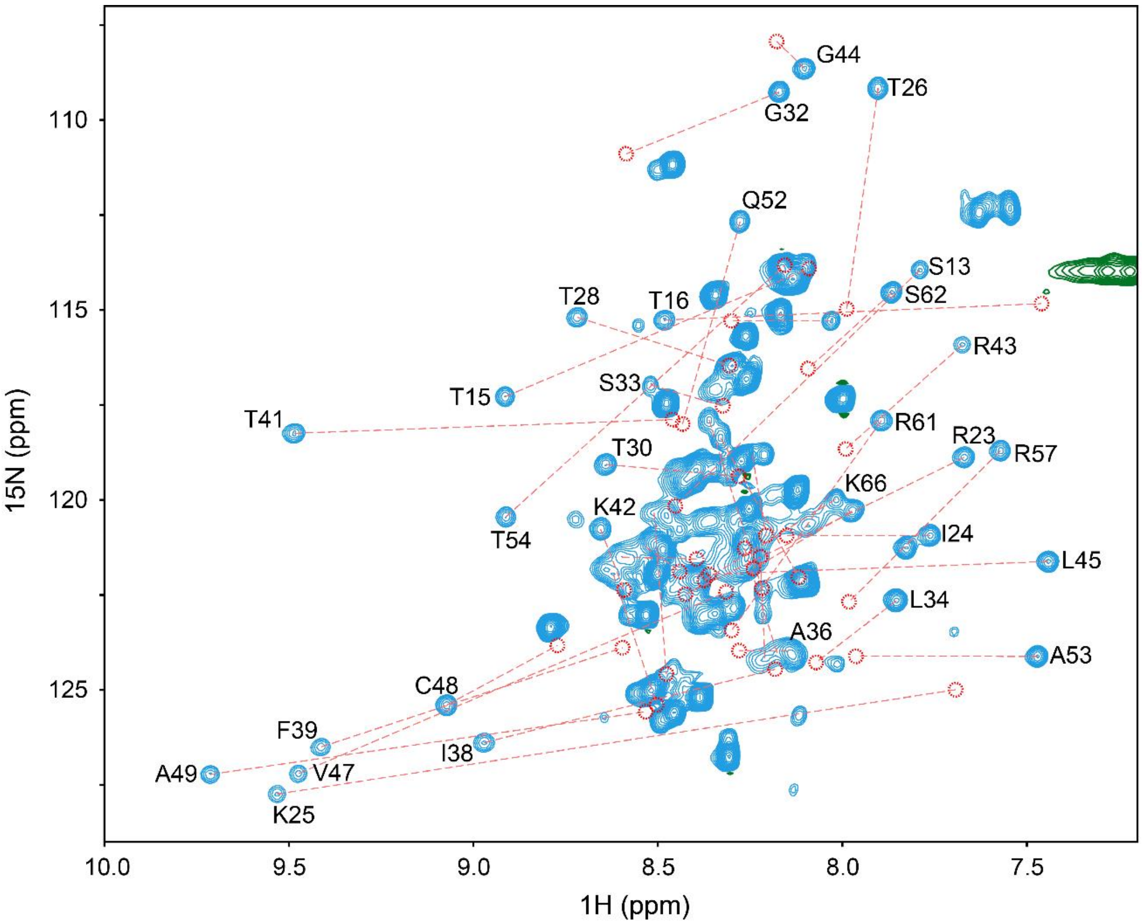


**Fig. S4. The ES is invisible in HSQC NMR spectra.** The reconstructed spectrum of the ES (red circles) overlaid on the Ltnαβ spectrum (blue). The red dashed lines connect the residue-specific chemical shifts of Ltnαβ and the ES.

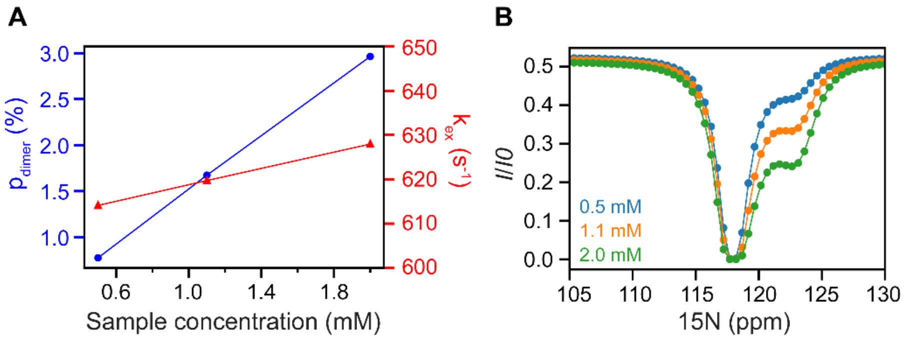


**Fig. S5. CEST profiles for symmetric dimerization are sensitive to sample concentration.** A) Variation in $p_E$ and $k_{ex}$ as a function of the total protein concentration calculated assuming the excited state to be a symmetric dimer (as described in Supplementary Text). B) CEST profiles simulated using the $p_E$ and $k_{ex}$ from panel (A).

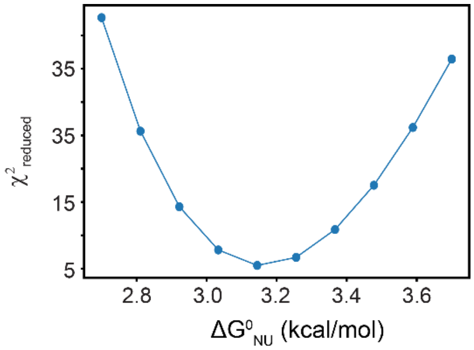


**Fig. S6. Analysis of urea unfolding of Ltnαβ at 20 °C.** The $\chi^2_{reduced}$ surface for $\Delta G^0_{NU}$ when the fluorescence-detected urea melt was fit by fixing the folding and unfolding baselines as described in detail in supplementary text.

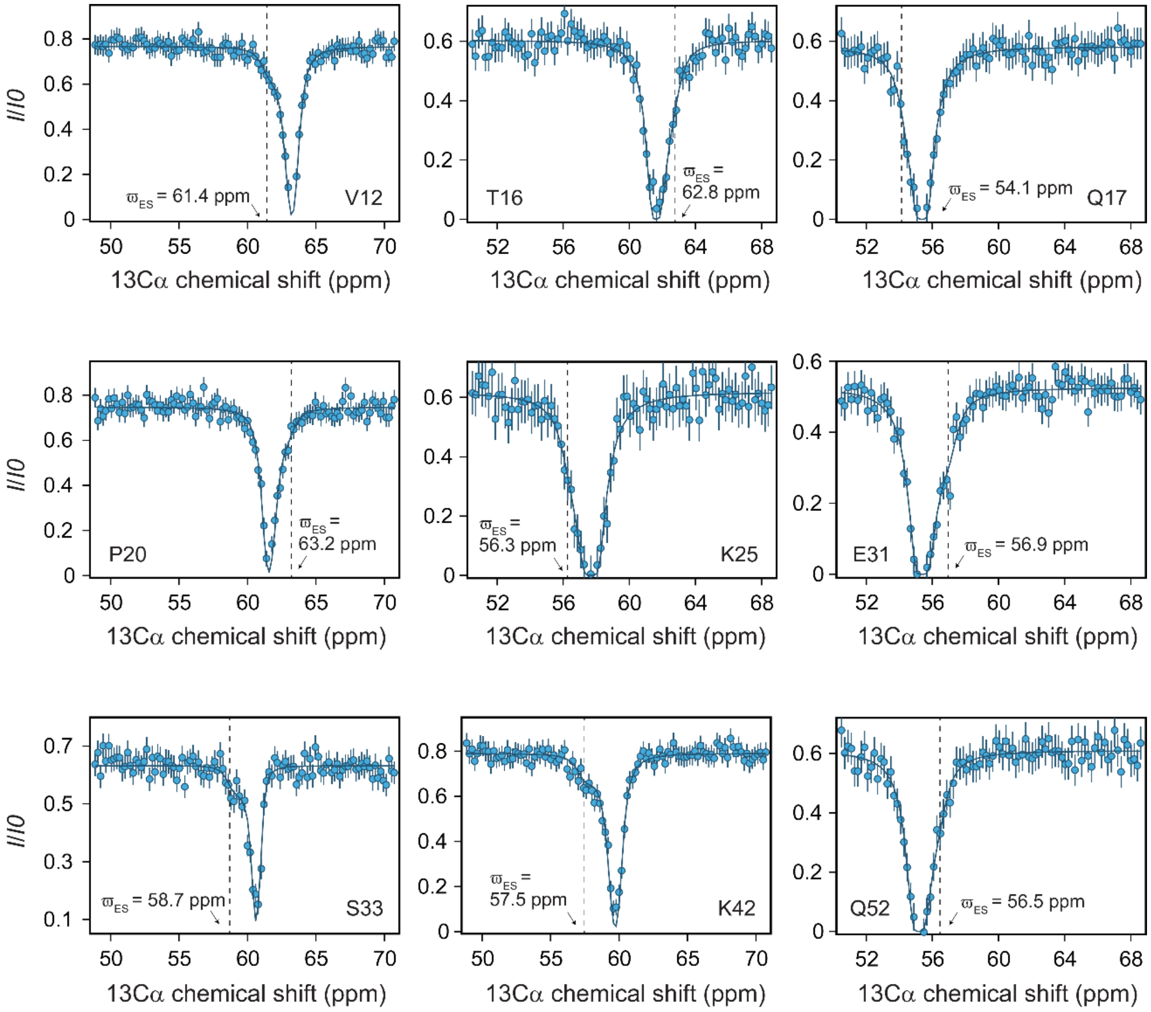


**Fig. S7. $^{13}$Cα CEST profiles reporting on Ltnαβ↔ES exchange**. $^{13}$Cα CEST profiles of V12, P20, S33 and K42 were recorded using a $^{1}$Hα→$^{13}$Cα (CEST)→$^{1}$Hα coherence transfer scheme[42]. CEST profiles of T16, Q17, K25, E31 and Q52 were acquired using an out-and-back coherence transfer scheme starting and ending on amide protons, chemical shift labelling $^{15}$N and carrying out CEST on $^{13}$Cα [41].

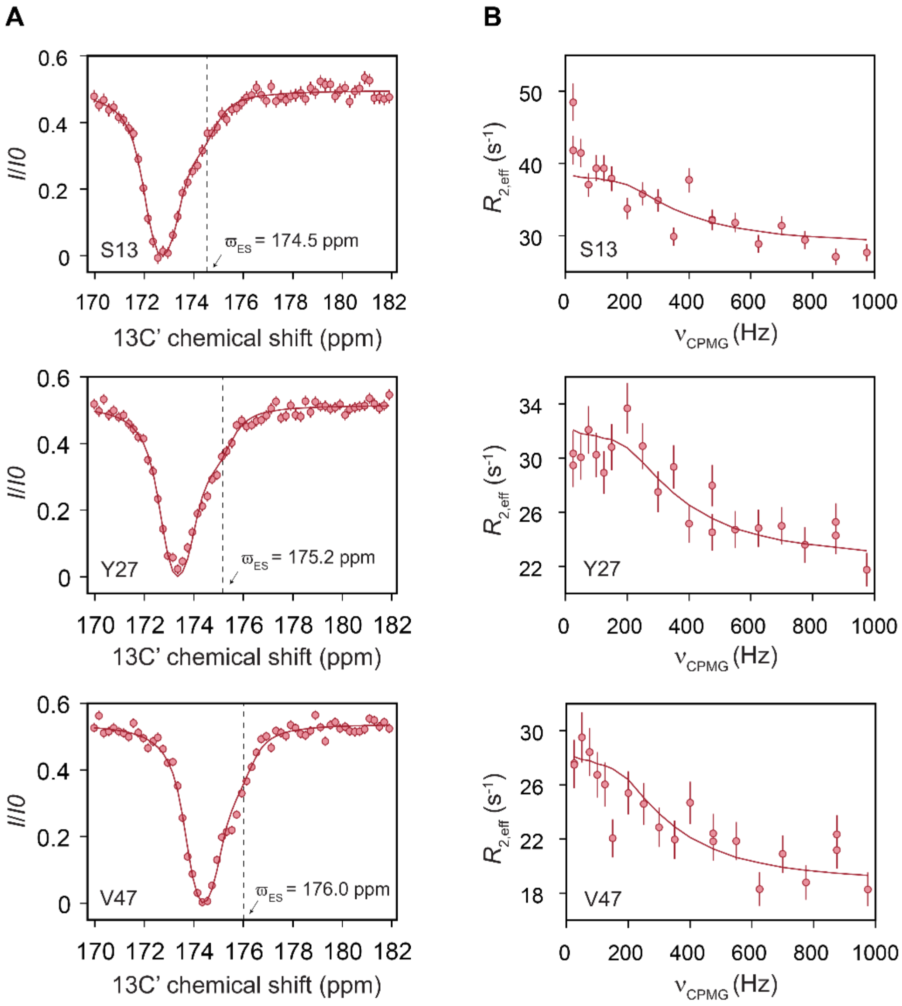


**Fig. S8. Detecting Ltnαβ↔ES exchange using the $^{13}$C' nucleus.** A) $^{13}$C' CEST profiles acquired using a $B_1$ field of 30 Hz. B) $^{13}$C' CPMG profiles acquired on a 700 MHz NMR spectrometer.

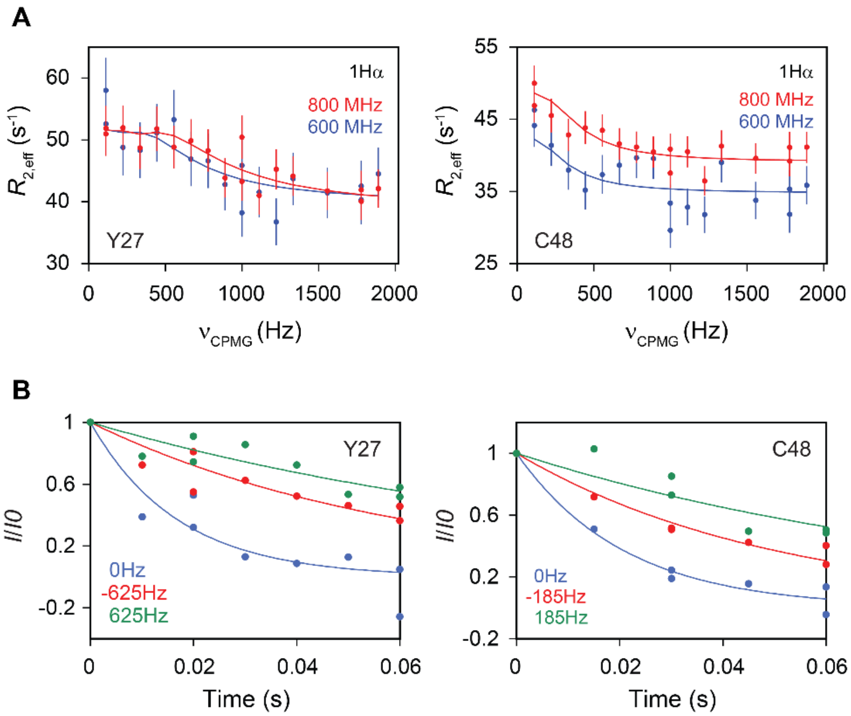


**Fig. S9. $^1$Hα CPMG profiles reporting on Ltnαβ↔ES exchange.** A) $^1$Hα CPMG profiles of Y27 and C48 acquired at a static magnetic field strength of 600 MHz (red) and 800 MHz (blue). B) $^1$Hα $R_{1\rho}$ profiles of Y27 and C48. The strengths and the offset positions of the weak spin-lock field were determined for each residue based on the Δϖ values obtained from the $^1$Hα CPMG profiles using the method described in[48] and are listed in Table S5.

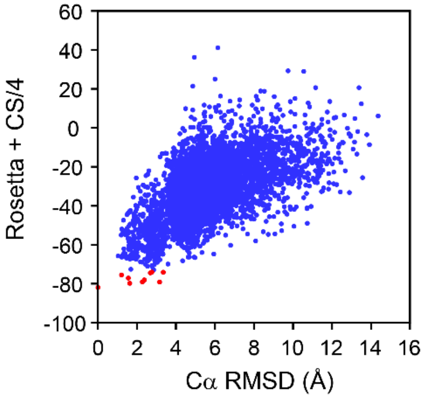


**Fig. S10. Calculating the structure of the ES.** The RMSD vs energy score for the CS-Rosetta structure calculation involving residues 9–52 of Ltn, where the energy of each conformation is calculated as the Rosetta energy added to one-fourth the chemical shift energy[56]. The plot is funnel-shaped, indicating an overall convergence of the structure calculation. A total of 5000 structures were generated during the calculation. The RMSD to the lowest energy structure was calculated using Cα nuclei for residues in the range 23–52. The 10 structures with the lowest energy are shown as red dots on the plot.

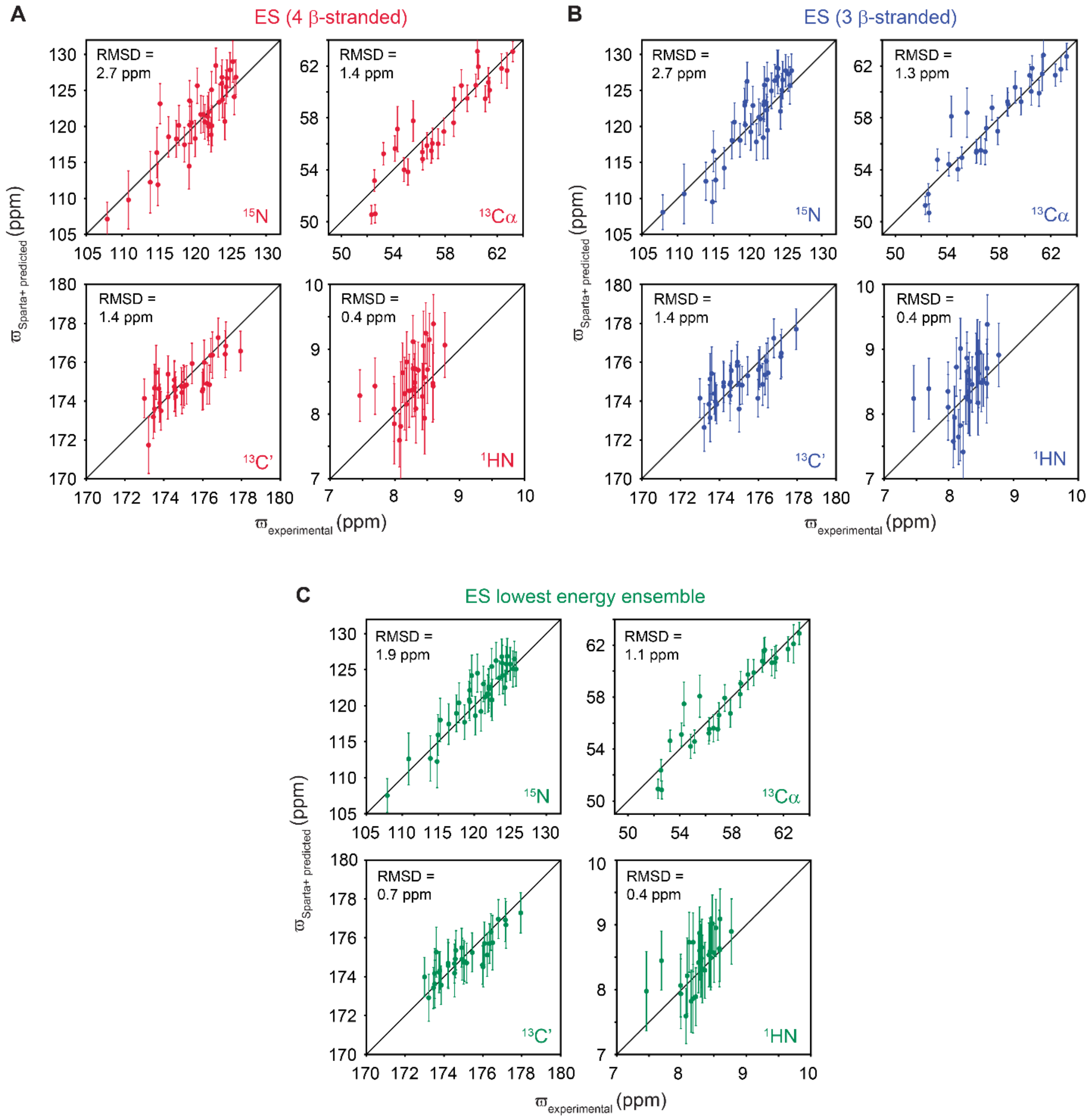


**Fig. S11. Predicted chemical shifts of the ES structural model agree well with input data.** A) $^{15}$N, $^{13}$Cα, $^{13}$C', and $^{1}$HN chemical shifts predicted using SPARTA+[59] for the 4-β-stranded and (B) the 3-β-stranded model of the ES plotted against the corresponding experimental chemical shifts. C) Correlation plots of chemical shifts averaged over the ensemble of the 10 lowest-energy ES structures plotted against the experimental chemical shifts. Errors in the averaged chemical shifts in panel C were calculated by propagating the prediction errors from individual structures. The RMSD values for each plot are mentioned on the plot which are within the range of prediction accuracy of SPARTA+ for each nuclei ($^{15}N$: 2.45 ppm, $^{13}C\alpha$:0.94 ppm, $^{13}C'$:1.09 ppm, and $^{1}HN$:0.49 ppm)[59].

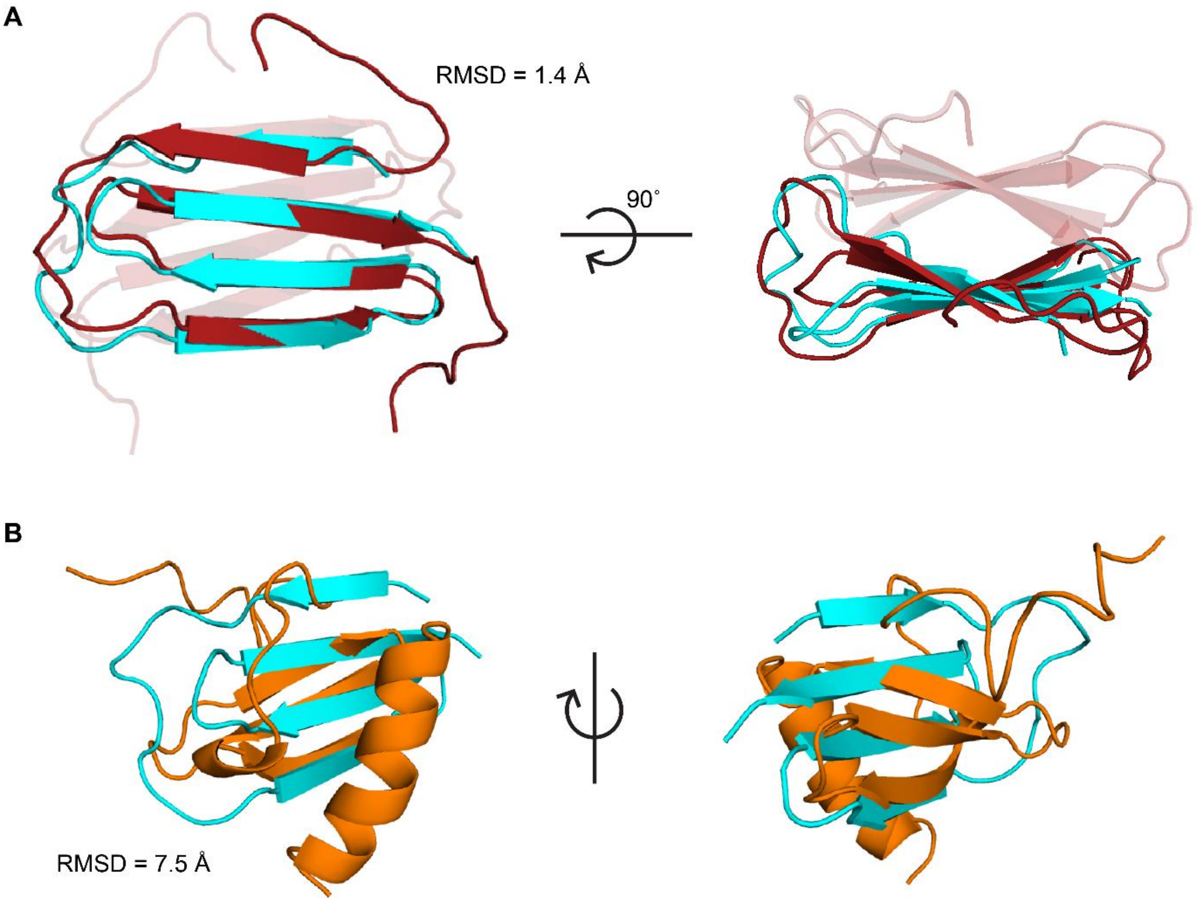


**Fig. S12. The β-sheet in the ES resembles the protomer of Ltnβ2 rather than the β-sheet in Ltnαβ.** A) Overlay of the ES (cyan) structure onto one protomer of Ltnβ2 (PDB:2JP1, red)[13] and (B) on Ltnαβ (PDB:1J9O, orange)[12]. Cα RMSD values are indicated for each of the overlays.

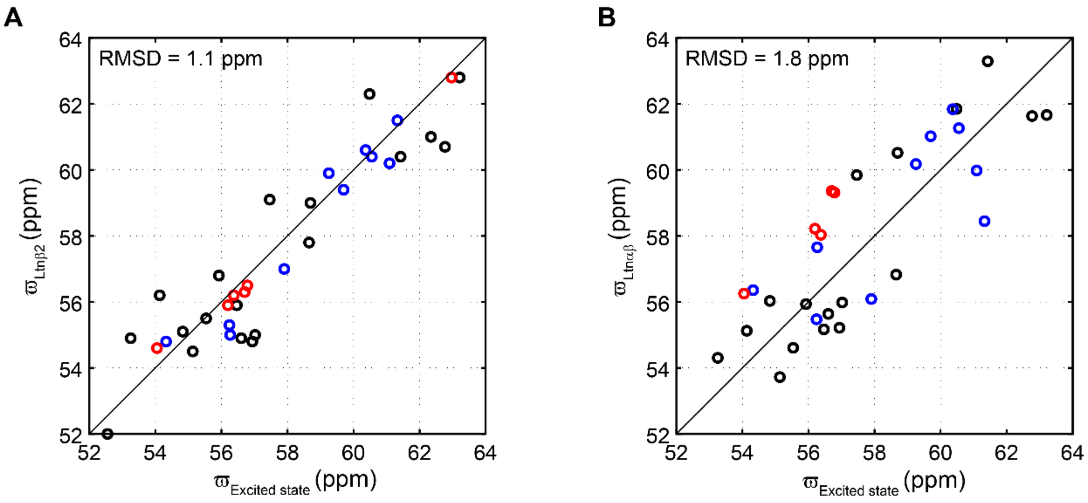


**Fig. S13. The $^{13}$Cα chemical shifts of the ES match better with those of Ltnβ2 than Ltnαβ.** A comparison of the ES $^{13}$Cα chemical shifts with those of Ltnβ2 (A) and Ltnαβ (B). Ltnβ2 chemical shifts were obtained from BMRB (accession ID:15215). Residues are color-coded according to their secondary structure in Ltnαβ: black circles indicate loop residues, red circles indicate residues in helical conformation, and blue circles indicate residues in β-strands. Solid lines in both panels depict the y=x function.

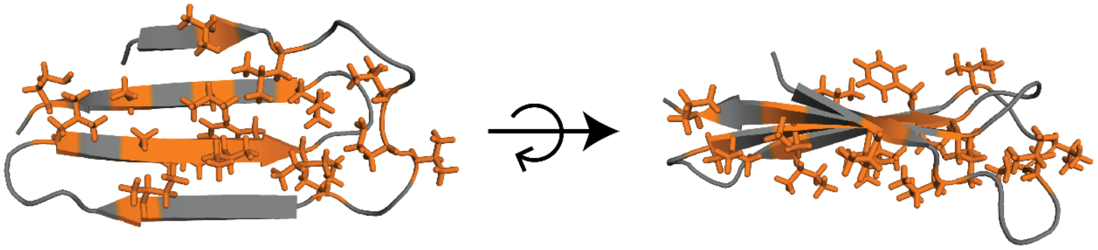

**Fig. S14. Hydrophobic residues in the ES are solvent exposed.** Hydrophobic residues (Ala, Phe, Leu, Met, Val, Ile, Pro) are highlighted in orange on the ES structure.

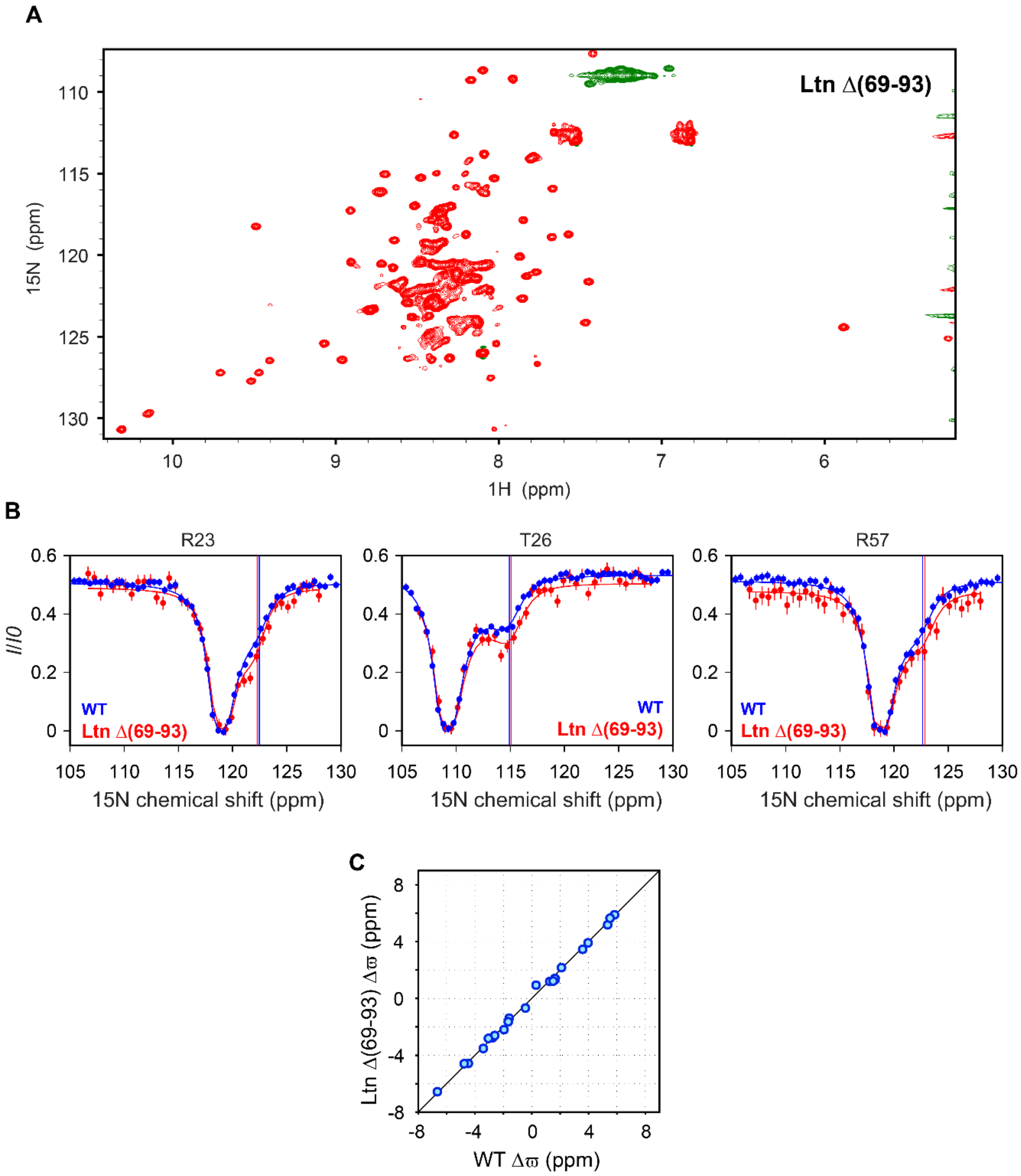


**Fig. S15. Truncating the disordered C-terminal tail (69–93) does not perturb the Ltnαβ↔ES equilibrium.** A) The $^{1}$H-$^{15}$N HSQC spectrum of LtnΔ(69–93). B) LtnΔ(69–93) (red) exhibits CEST profiles very similar to those of the WT protein (blue). C) A comparison of the chemical shift differences between the ES and Ltnαβ for LtnΔ(69–93) (y-axis) and WT (x-axis) Ltn. The solid line depicts the y=x function. The correlation between the two parameters is good, indicating that LtnΔ(69–93) samples the same ES conformation as WT.

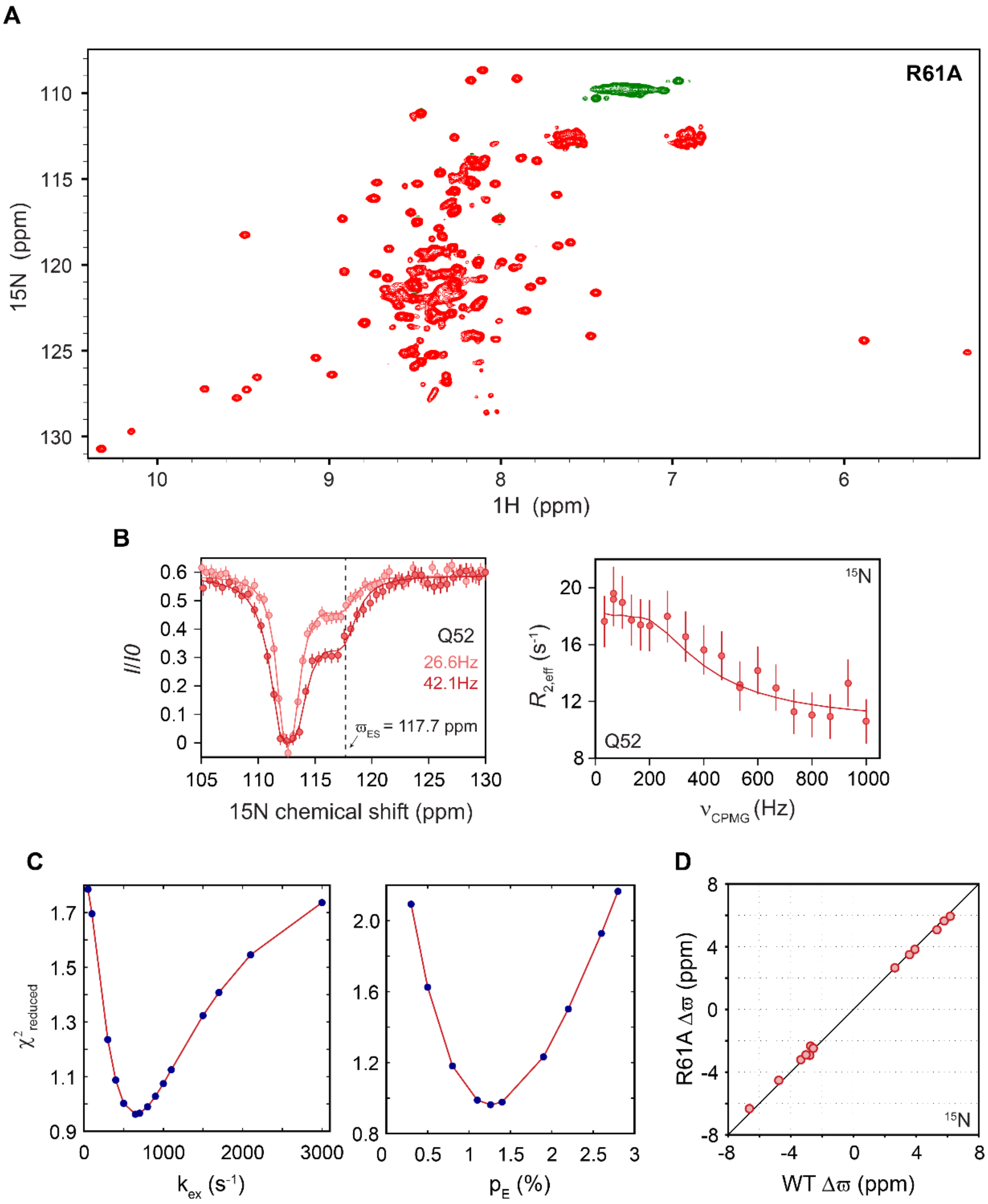


**Fig. S16. Characterization of Ltnαβ↔ES exchange in the R61A mutant of Ltn.** A) The $^{1}$H-$^{15}$N HSQC spectrum of R61A Ltn. B) $^{15}$N CEST (left) and CPMG (right) profiles for the residue Q52, showing the presence of an excited state. The dashed line in the CEST profile indicates the chemical shift of the ES. C) Prominent minima observed in the $\chi^2_{reduced}$ surfaces of $k_{ex}$ (left) and $p_E$ (right) indicates that these parameters can be obtained reliably from fitting the CEST and CPMG data. D) A comparison of the chemical shift differences between the ES and Ltnαβ for R61A (y-axis) and WT (x-axis) Ltn. The solid line depicts the y=x function. The correlation between the two parameters is good, indicating that R61A Ltn samples the same ES conformation as WT.

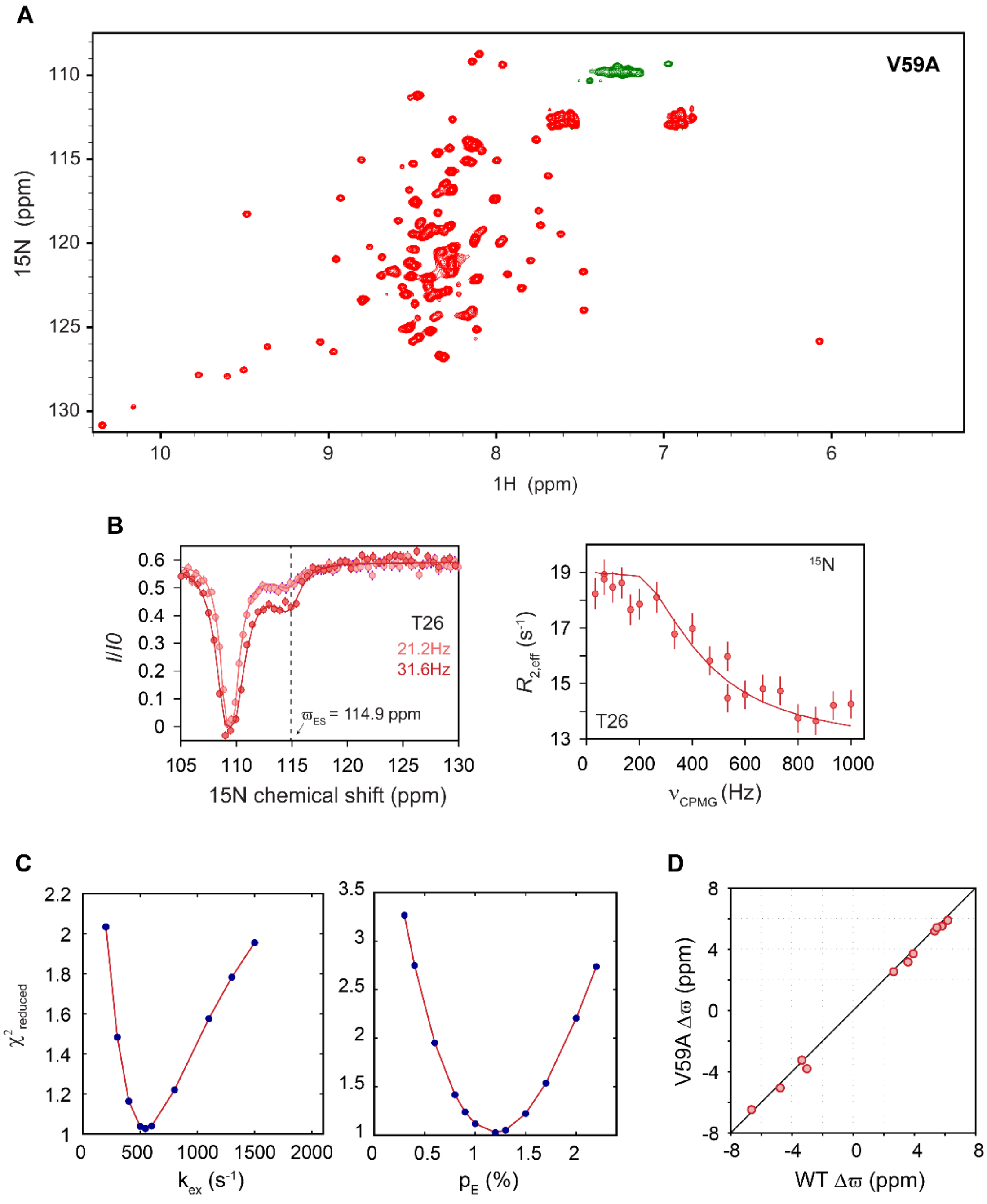


**Fig. S17. Characterization of Ltnαβ↔ES exchange in the V59A mutant of Ltn.** A) The $^{1}$H-$^{15}$N HSQC spectrum of the V59A mutant. B) $^{15}$N CEST (left) and CPMG (right) profiles for the residue T26, showing the presence of an excited state. The dashed line in the CEST profile indicates the chemical shift of the ES. C) Prominent minima observed in the $\chi^2_{reduced}$ surfaces of $k_{ex}$ (left) and $p_E$ (right) indicates that these parameters can be obtained reliably from fitting the CEST and CPMG data. D) A comparison of the chemical shift differences between the ES and

Ltnαβ for V59A (y-axis) and WT (x-axis) Ltn. The solid line depicts the y=x function. The correlation between the two parameters is good, indicating that V59A Ltn samples the same ES conformation as WT.

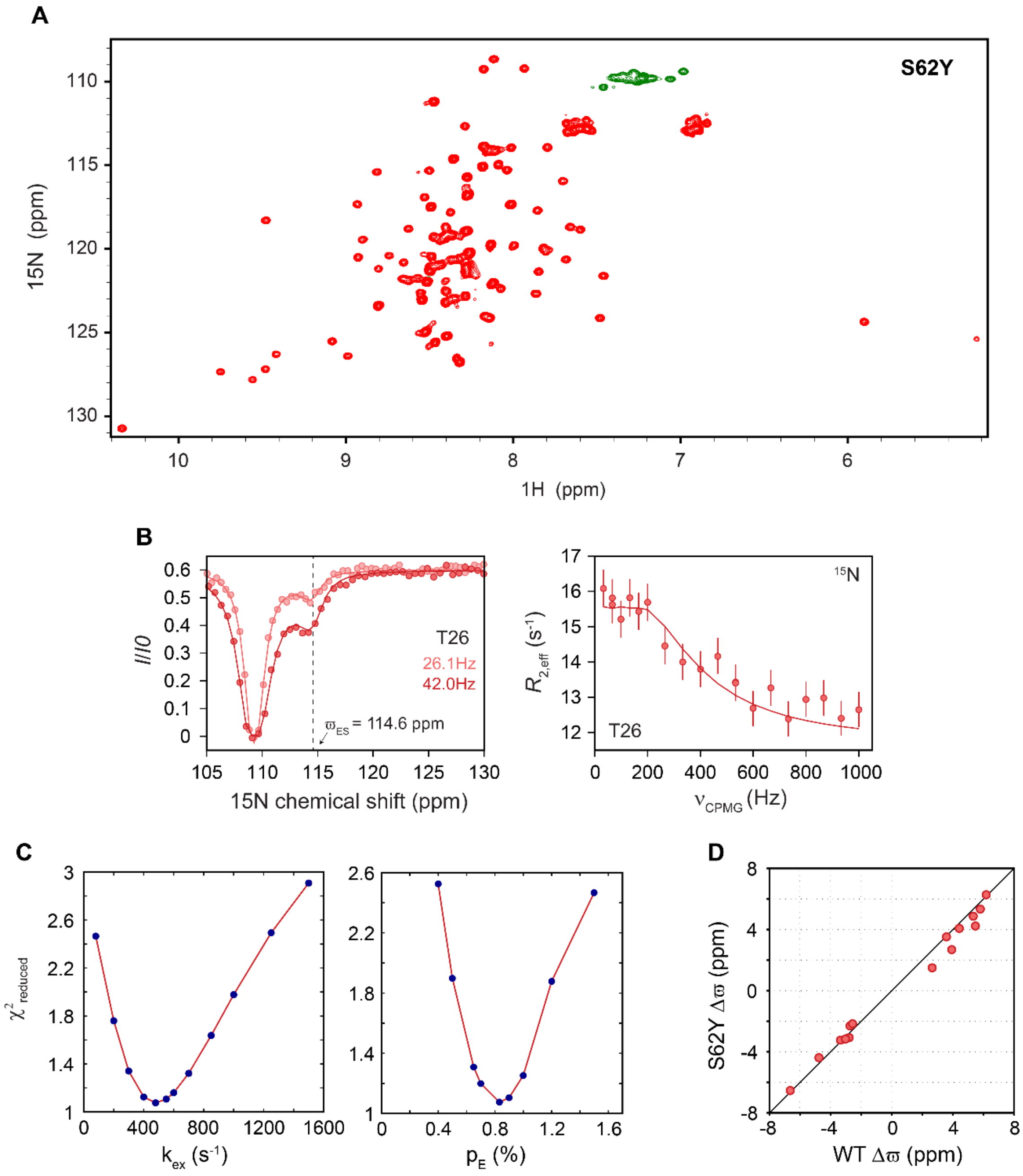


**Fig. S18. Characterization of Ltnαβ↔ES exchange in the S62Y mutant of Ltn .** A) The $^{1}$H-$^{15}$N HSQC spectrum of S62Y Ltn. B) $^{15}$N CEST (left) and CPMG (right) profiles for the residue T26, showing the presence of an excited state. The dashed line in the CEST profile indicates the chemical shift of the ES. C) Prominent minima observed in the $\chi^2_{reduced}$ surfaces of $k_{ex}$ (left) and $p_E$ (right) indicates that these parameters can be obtained reliably from fitting the CEST and CPMG data. D) A comparison of the chemical shift differences between the ES and Ltnαβ for S62Y (y-axis) and WT (x-axis) Ltn. The solid line depicts the y=x function. The correlation between the two parameters is good, indicating that S62Y Ltn samples the same ES conformation as WT.

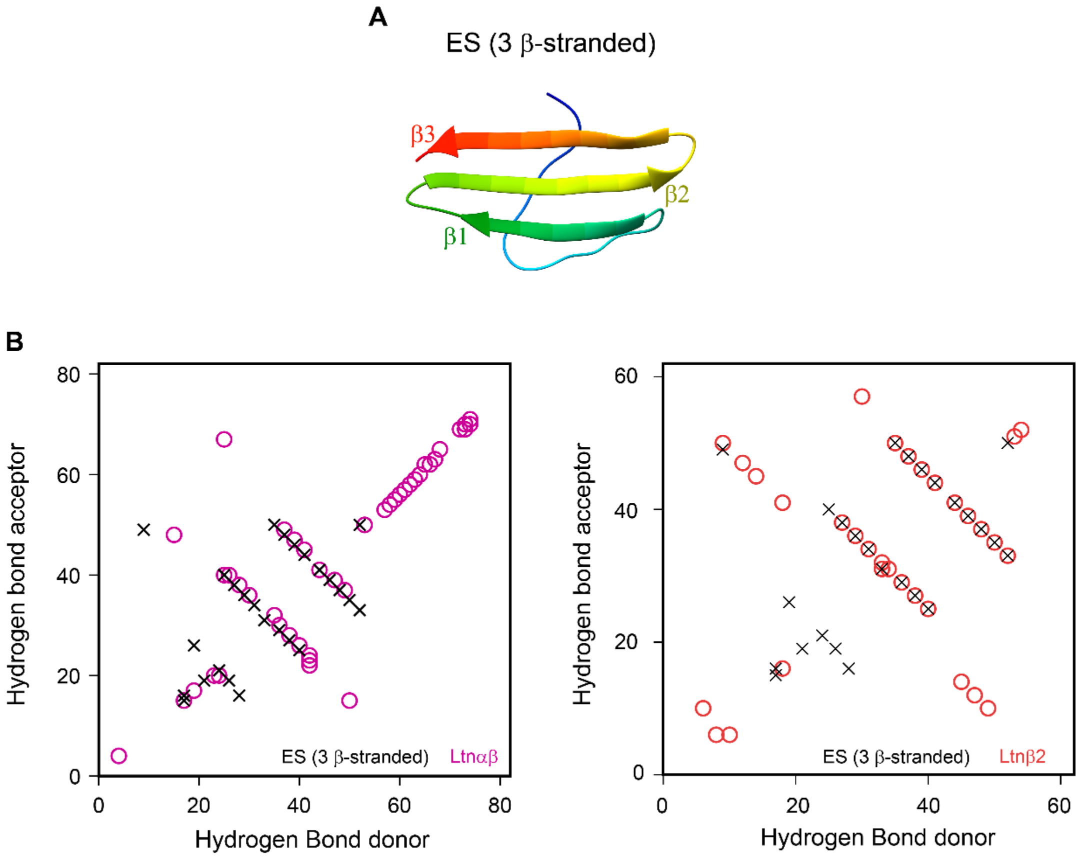


**Fig. S19. Characterizing the hydrogen bonding network in the 3-β-stranded ES model.** A) Representative 3-β-stranded model of the ES. B) Comparison of the hydrogen bonds present in the ES (3-β-stranded, black cross) and with those in Ltnαβ (left, pink circles) and Ltnβ2 (right, orange circles). The hydrogen bond donor residue is plotted on the x-axis and the acceptor is plotted on the y-axis.

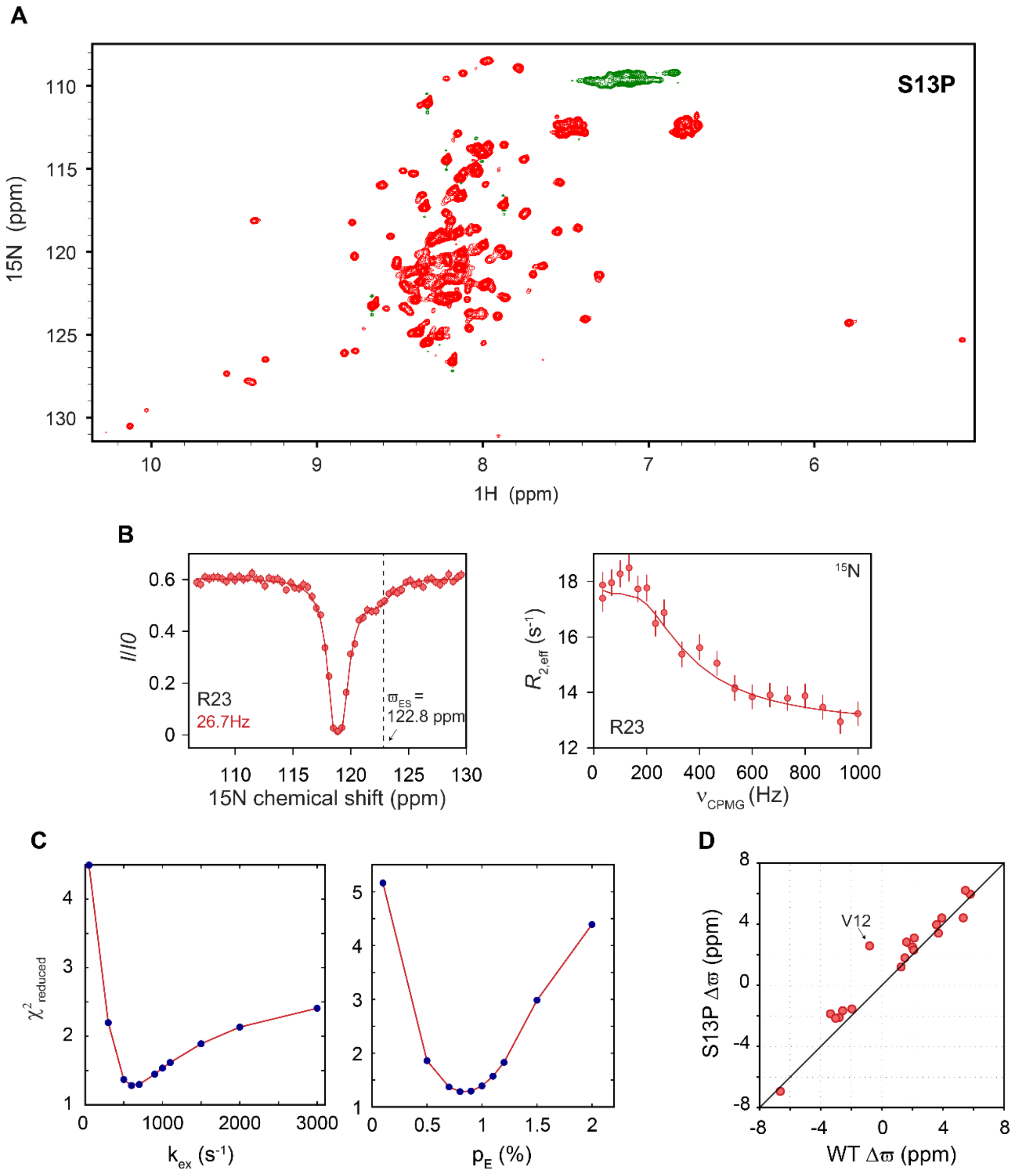


**Fig. S20. Characterization of Ltnαβ↔ES exchange in the S13P mutant of Ltn.** A) The $^1$H-$^{15}$N HSQC spectrum of the S13P mutant. B) $^{15}$N CEST (left) and CPMG (right) profiles for the residue R23, showing the presence of an excited state. The dashed line in the CEST profile indicates the chemical shift of the ES. C) Prominent minima observed in the $\chi^2_{reduced}$ surfaces of $k_{ex}$ (left) and $p_E$ (right) indicates that these parameters can be obtained reliably from fitting the CEST and CPMG data. D) A comparison of the chemical shift differences between the ES and Ltnαβ for S13P (y-axis) and WT (x-axis) Ltn. The solid line depicts the y=x function. The correlation between the two parameters is good, indicating that S13P Ltn samples the same ES conformation as WT.

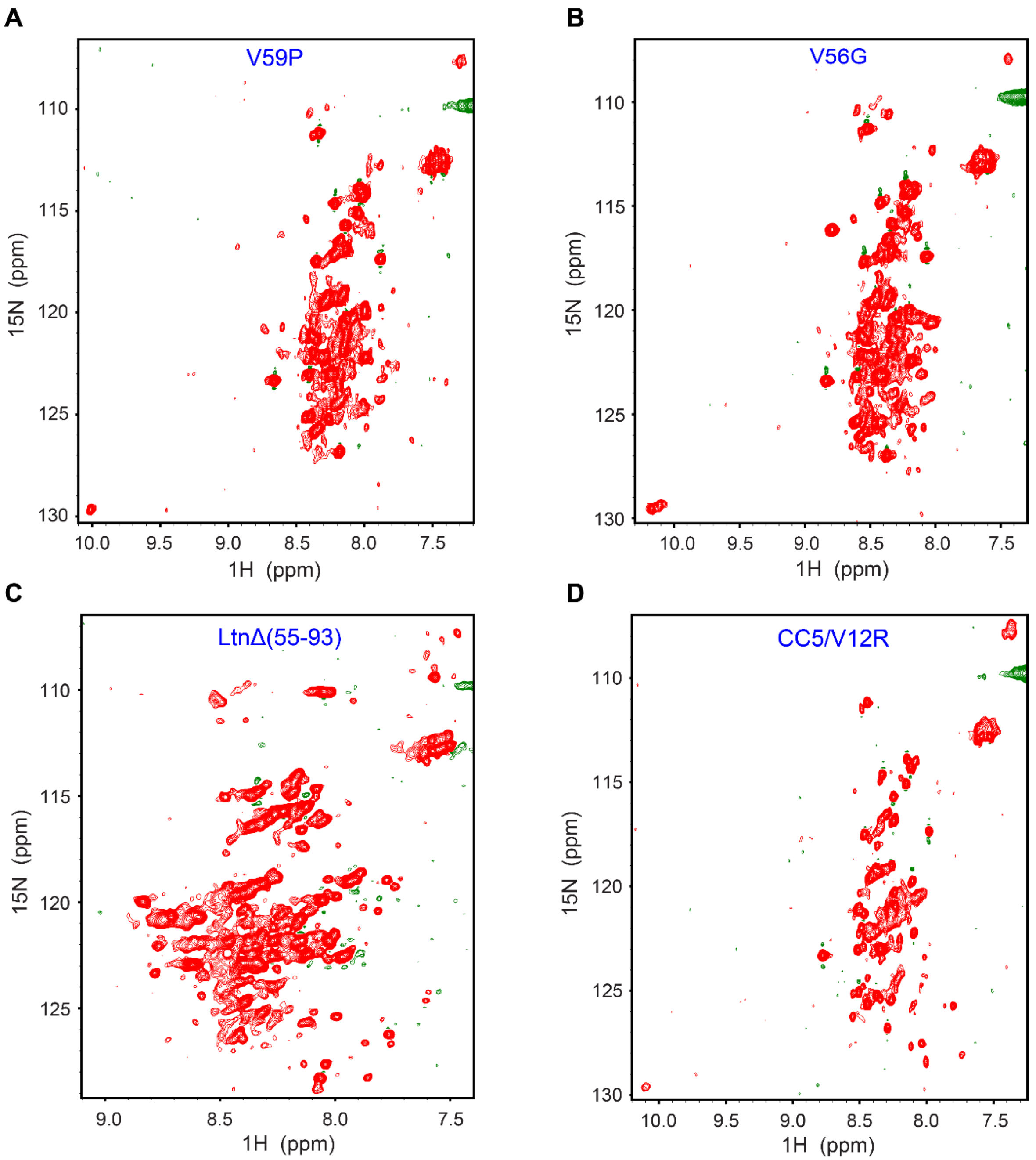


**Fig. S21. Engineering mutations to stabilize the ES of WT Ltn.** $^{1}$H-$^{15}$N HSQC spectra of V59P (A), V56G (B), LtnΔ55–93 (C), and CC5/V12R (D) Ltn designed to selectively stabilize the ES conformation.

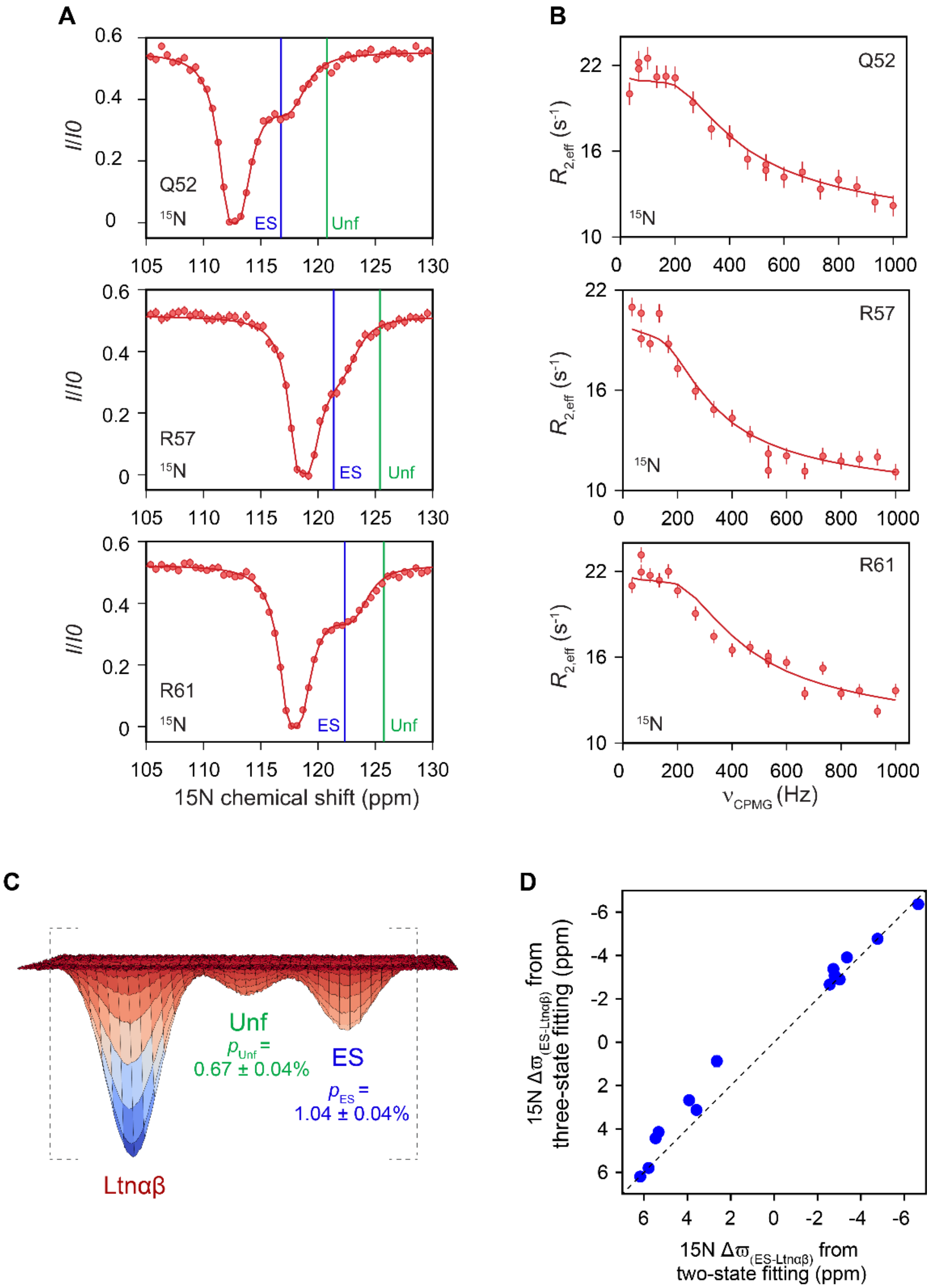


**Fig. S22. Modeling the $^{15}$N CEST and CPMG profiles of WT Ltn to the three-state Ltnαβ↔Unf↔ES scheme.** A) CEST and B) CPMG profiles of Q52, R57 and R61. Solid red lines are fits to the three-state model Ltnαβ↔Unf↔ES. Vertical lines indicate the chemical shift positions of the ES (blue line) and the unfolded state (Unf, green line). B) Schematic free energy surface indicating the relative populations of the unfolded state (green) and the ES (blue) obtained from three-state modelling of the CEST and CPMG data. C) Comparison of the $^{15}$N $\Delta\varpi$ values for the Ltnαβ-ES transition obtained from two-state fitting (x-axis) and three-state fitting (y-axis), demonstrating that the ES chemical shifts are largely independent of the fitting model. The dashed line depicts the y=x function.

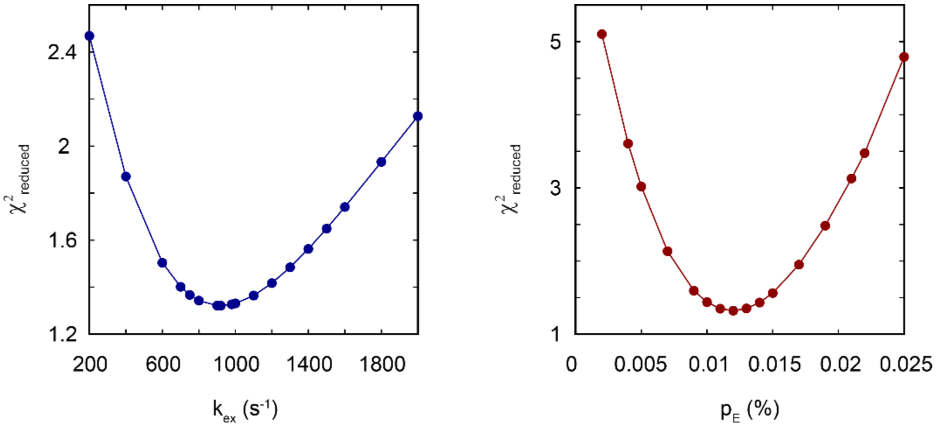


**Fig. S23. $\chi^2_{reduced}$ surfaces for the $^{13}$Cα CEST data.** $\chi^2_{reduced}$ surfaces for $k_{ex}$ (left) and $p_E$ (right) obtained by fitting the $^{13}$Cα CEST profiles of V59.

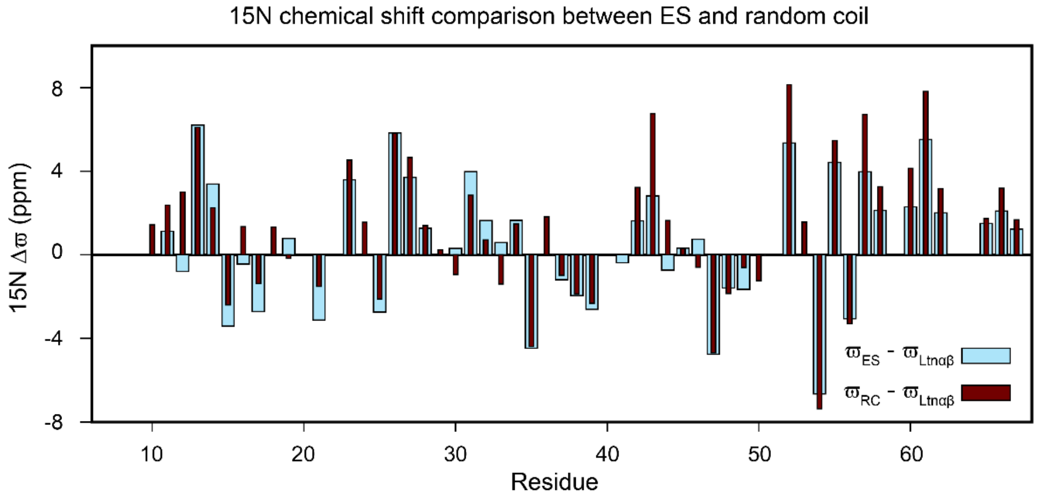
15N chemical shift comparison between ES and random coil
15N Δϖ (ppm)
Residue
ϖES - ϖLtnαβ
ϖRC - ϖLtnαβ

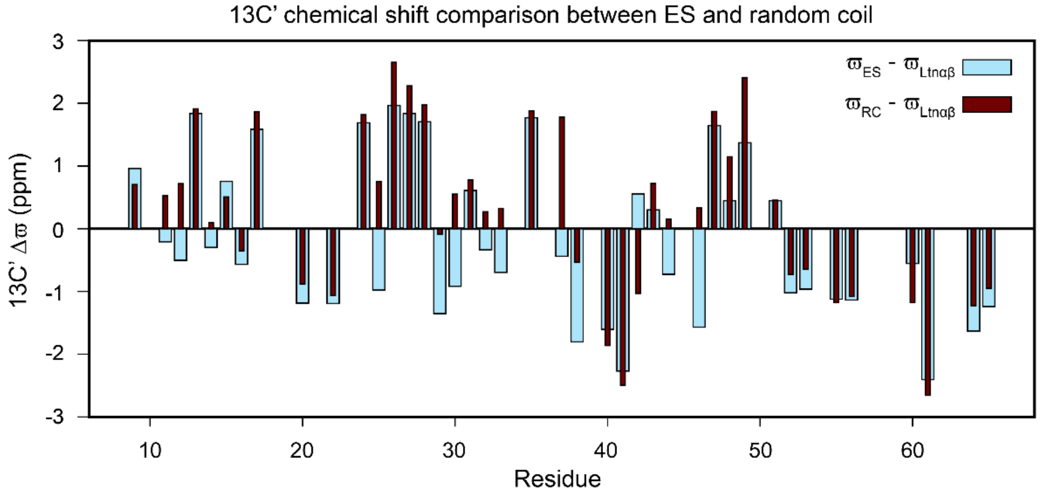
13C' chemical shift comparison between ES and random coil
13C' Δϖ (ppm)
Residue
ϖES - ϖLtnαβ
ϖRC - ϖLtnαβ

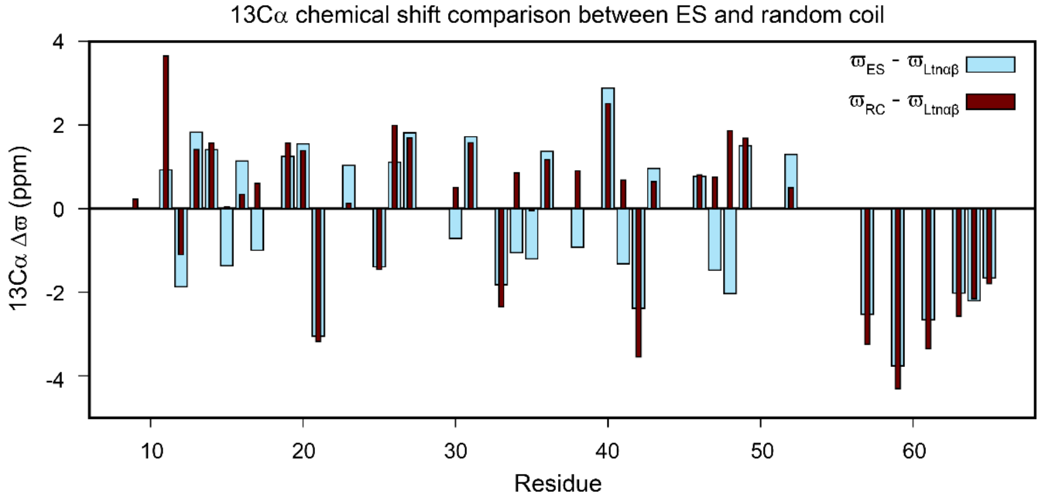
13Cα chemical shift comparison between ES and random coil
13Cα Δϖ (ppm)
Residue
ϖES - ϖLtnαβ
ϖRC - ϖLtnαβ

**Fig. S24. Chemical shift comparison between the ES and random coil.** Bar plots comparing the $^{15}$N (top), $^{13}$C' (middle), and $^{13}$Cα (bottom) (ES-Ltnαβ) chemical shift differences (light blue) with (RC-Ltnαβ) (brown).

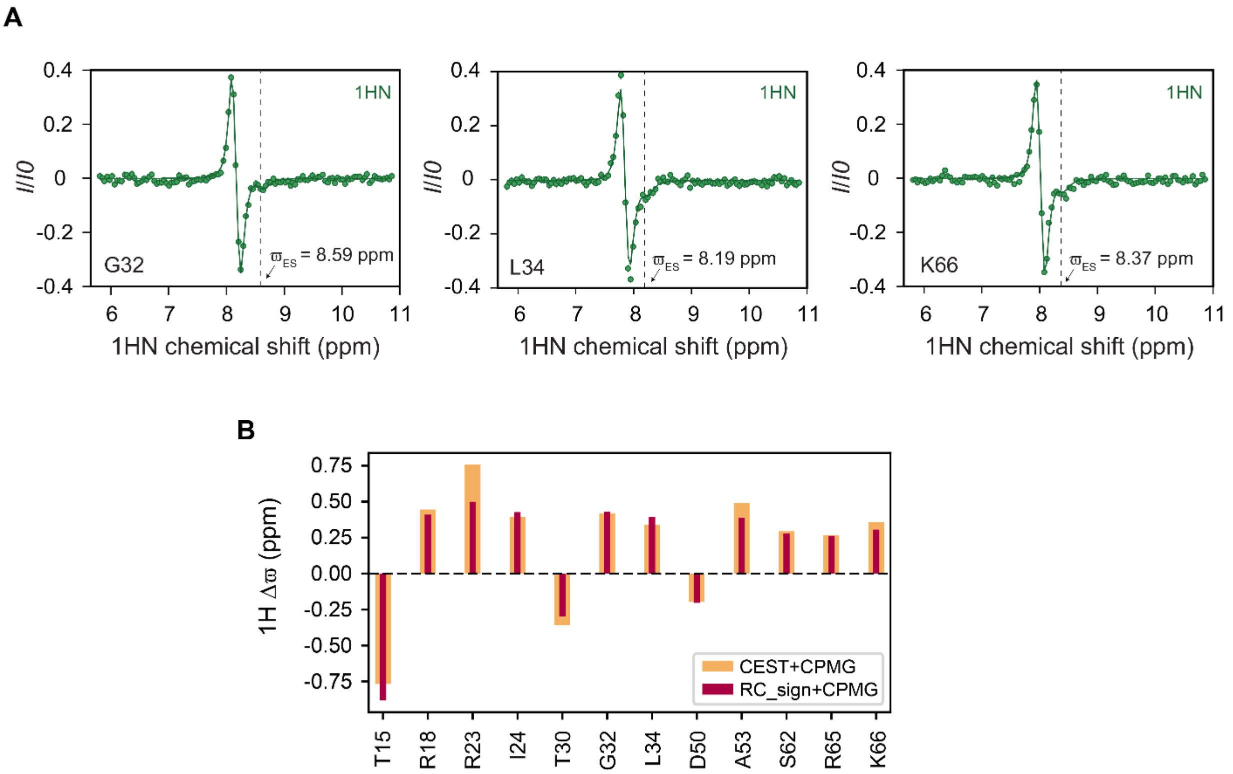


**Fig. S25. The Ltnαβ-ES exchange viewed through the amide proton nucleus.** A) $^{1}$HN CEST profiles of G32, L34 and K66. Dashed lines indicate the chemical shift positions of the ES. B) Bar plot comparing the $\Delta\varpi$ values obtained from global fitting of $^{1}$H CEST and CPMG (orange bars) against the $|\Delta\varpi|$ obtained from CPMG (red bars), where the signs for $\Delta\varpi$ are chosen to be the same as for the difference between the random coil (RC) conformation and Ltnαβ.

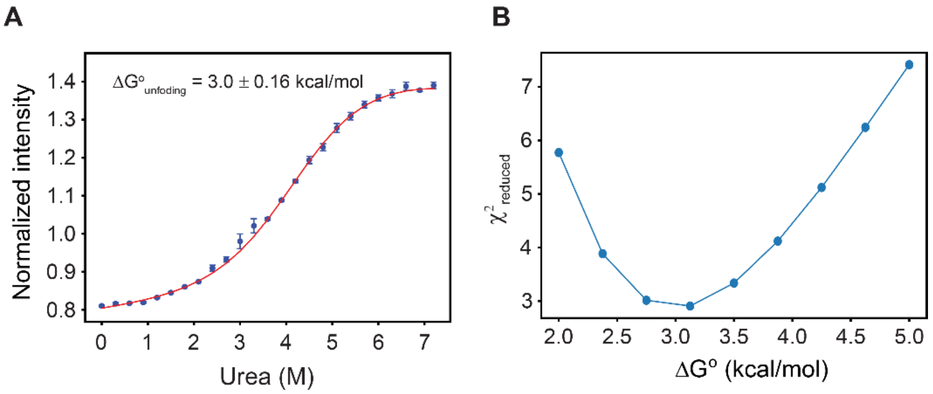


**Fig. S26. Analysis of the urea unfolding of Ltnαβ at 20 °C.** A) Normalized fluorescence intensity of Ltnαβ as a function of the urea concentration. Solid lines are fits of the unfolding profile obtained without fixing the folding and unfolding baselines. B) $\chi^2_{reduced}$ surface for $\Delta G^0$ of unfolding obtained using this fitting method.

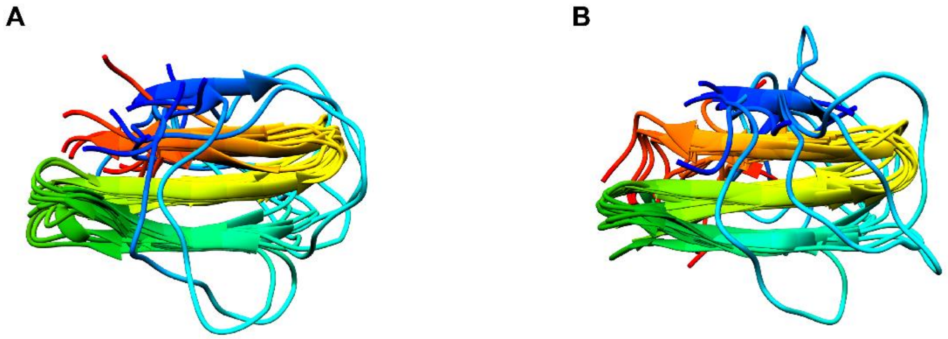


**Fig. S27. Variations in the structural ensemble of the ES on the input and calculation protocol.** A) Ensemble of the 10 lowest-energy structures calculated for 9–52 including the $^{1}$Hα chemical shifts. A total of 5,000 structures were generated. B) Ensemble of the 10 lowest-energy structures calculated for 9–54 without $^{1}$Hα chemical shifts. A total of 50,000 structures were generated.

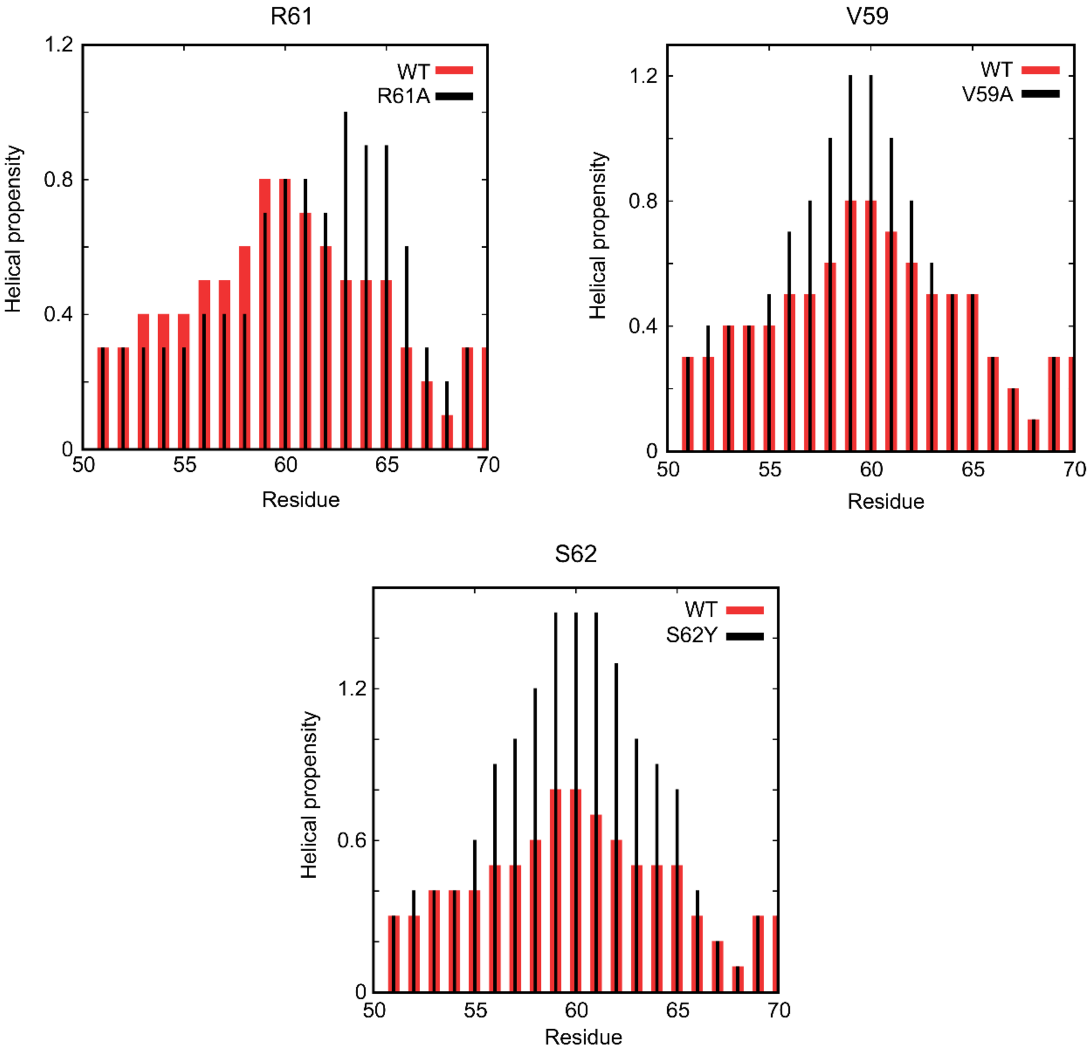


**Fig. S28. Helical propensities of the engineered mutants of Ltn.** Predicted helical propensity for the C-terminal helical region of Ltn mutants (black lines) compared with wild-type (WT) Ltn (red bars).

**Table S1:** List of chemical shifts for the excited state of WT Ltn.

| Residue | Nuclei | $\Delta\varpi$ (ppm) | error (ppm) | $\varpi_{ES}$ (ppm) |
|---|---|---|---|---|
| R9 | CA | -0.0109 | 15.5151 | 55.925 |
| R9 | CO | 0.9611 | 0.0693 | 176.889 |
| T10 | N | -0.0009 | 0.5393 | 115.283 |
| T10 | H | 0.2696 | 0.0195 | 8.301 |
| C11 | N | 1.1215 | 0.1031 | 120.444 |
| C11 | CA | 0.9266 | 0.3923 | 55.536 |
| C11 | CO | -0.2081 | 0.2081 | 173.518 |
| V12 | N | -0.7991 | - | 119.647 |
| V12 | CA | -1.8662 | 0.107 | 61.425 |
| V12 | CO | -0.5065 | 0.475 | 174.931 |
| S13 | N | 6.2034 | 0.0766 | 120.157 |
| S13 | CA | 1.8281 | 0.2191 | 58.655 |
| S13 | CO | 1.8359 | 0.106 | 174.543 |
| S13 | H | 0.7273 | 0.0295 | 8.453 |
| L14 | N | 3.3852 | 0.0628 | 125.392 |
| L14 | CA | 1.41 | 0.234 | 55.133 |
| L14 | CO | -0.2976 | 0.7054 | 177.157 |
| L14 | H | -0.108 | 0.0263 | 8.503 |
| T15 | N | -3.4128 | 0.0572 | 113.911 |
| T15 | CA | -1.3628 | 0.1811 | 60.485 |
| T15 | CO | 0.756 | 0.0729 | 175.012 |
| T15 | H | -0.7675 | 0.0414 | 8.09 |
| T16 | N | -0.4472 | 0.2624 | 114.832 |
| T16 | CA | 1.1383 | 0.1313 | 62.77 |
| T16 | CO | -0.5694 | 0.1156 | 174.218 |
| T16 | H | -1.0209 | 0.1071 | 7.46 |
| Q17 | N | -2.7178 | 0.0604 | 121.496 |
| Q17 | CA | -0.9953 | 0.1781 | 54.131 |
| Q17 | CO | 1.5864 | 0.0607 | 175.453 |
| Q17 | H | 0.0947 | 0.0173 | 8.222 |
| R18 | N | 0.0027 | 0.8162 | 121.273 |
| R18 | H | 0.4452 | 0.0215 | 8.265 |
| L19 | N | 0.7812 | 0.154 | 125.817 |
| L19 | CA | 1.2487 | 0.1641 | 52.551 |
| P20 | CA | 1.5499 | 0.1724 | 63.213 |
| P20 | CO | -1.1856 | 0.1519 | 176.502 |

| | | | | |
|---|---|---|---|---|
| V21 | N | -3.1229 | 0.0676 | 119.309 |
| V21 | CA | -3.0488 | 0.1548 | 62.345 |
| V21 | HA | 0.2878 | 0.0351 | 3.905844 |
| S22 | CO | -1.1925 | 0.1125 | 174.209 |
| R23 | N | 3.586 | 0.0532 | 122.48 |
| R23 | CA | 1.0368 | 0.1764 | 57.02 |
| R23 | H | 0.7567 | 0.0262 | 8.425 |
| I24 | N | 0.0017 | 0.5726 | 120.935 |
| I24 | CO | 1.6868 | 0.1311 | 176.078 |
| I24 | H | 0.3924 | 0.0219 | 8.151 |
| K25 | N | -2.7474 | 0.0791 | 124.996 |
| K25 | CA | -1.3899 | 0.1993 | 56.266 |
| K25 | CO | -0.9781 | 0.0888 | 174.91 |
| K25 | H | -1.8373 | 0.222 | 7.693 |
| T26 | N | 5.8319 | 0.0466 | 114.972 |
| T26 | CA | 1.1102 | 0.1634 | 61.092 |
| T26 | CO | 1.9625 | 0.0896 | 173.459 |
| T26 | H | 0.0821 | 0.0238 | 7.988 |
| Y27 | N | 3.7047 | 0.0395 | 122.028 |
| Y27 | CA | 1.8129 | 0.3461 | 57.902 |
| Y27 | CO | 1.8373 | 0.0824 | 175.167 |
| Y27 | H | -0.2061 | 0.0225 | 8.119 |
| Y27 | HA | -1.1349 | 0.1491 | 4.80814 |
| T28 | N | 1.2645 | - | 116.46 |
| T28 | CO | 1.7056 | 0.1701 | 173.855 |
| T28 | H | -0.4194 | 0.0442 | 8.307 |
| I29 | N | -0.0119 | 4.3692 | 123.023 |
| I29 | CO | -1.3531 | 0.0723 | 175.044 |
| T30 | N | 0.3036 | 0.3286 | 119.378 |
| T30 | CA | -0.7134 | 0.2811 | 60.553 |
| T30 | CO | -0.9183 | 0.1139 | 172.998 |
| T30 | H | -0.3598 | 0.0259 | 8.281 |
| E31 | N | 3.9781 | 0.1261 | 124.581 |
| E31 | CA | 1.7212 | 0.1142 | 56.937 |
| E31 | CO | 0.6085 | 0.0485 | 176.794 |
| E31 | H | -0.0002 | 0.1522 | 8.477 |
| G32 | N | 1.6409 | 0.0707 | 110.89 |
| G32 | CO | -0.3346 | 0.0604 | 173.599 |

| | | | | |
|---|---|---|---|---|
| G32 | H | 0.4191 | 0.0144 | 8.586 |
| S33 | N | 0.5883 | 0.2928 | 117.523 |
| S33 | CA | -1.8211 | 0.1808 | 58.696 |
| S33 | CO | -0.6961 | 0.0506 | 173.788 |
| S33 | H | -0.1952 | 0.0355 | 8.324 |
| L34 | N | 1.6465 | 0.0526 | 124.278 |
| L34 | CA | -1.0628 | 0.1316 | 53.253 |
| L34 | H | 0.3384 | 0.0122 | 8.071 |
| R35 | N | -4.4713 | 0.0665 | 122.037 |
| R35 | CA | -1.2003 | 0.0781 | 54.83 |
| R35 | CO | 1.7688 | 0.0907 | 175.979 |
| A36 | N | 0.0054 | 2.0434 | 123.964 |
| A36 | CA | 1.3695 | 0.1858 | 52.607 |
| A36 | H | 0.1369 | 0.0153 | 8.28 |
| V37 | N | -1.1938 | 0.0807 | 119.385 |
| V37 | CO | -0.4402 | 0.1807 | 173.748 |
| I38 | N | -1.9546 | 0.0757 | 124.455 |
| I38 | CA | -0.9248 | 0.212 | 59.252 |
| I38 | CO | -1.8039 | 0.1028 | 174.535 |
| I38 | H | -0.7856 | 0.1218 | 8.182 |
| F39 | N | -2.6075 | 0.0653 | 123.889 |
| F39 | H | -0.8155 | 0.1556 | 8.596 |
| F39 | HA | -0.5344 | 0.1137 | 4.826576 |
| I40 | CA | 2.8821 | 0.1985 | 61.33 |
| I40 | CO | -2.1799 | 0.1314 | 176.349 |
| I40 | HA | -0.6 | 0.1321 | 4.675974 |
| T41 | N | -0.3838 | 0.2767 | 117.883 |
| T41 | CA | -1.3206 | 0.1212 | 59.699 |
| T41 | CO | -2.269 | 0.1095 | 174.612 |
| T41 | H | -1.0228 | 0.102 | 8.46 |
| K42 | N | 1.6217 | 0.0723 | 122.371 |
| K42 | CA | -2.3837 | 0.12 | 57.461 |
| K42 | CO | 0.5546 | 0.1497 | 177.949 |
| K42 | H | 0.0641 | 0.052 | 8.592 |
| K42 | HA | 0.2969 | 0.042 | 4.253881 |
| R43 | N | 2.8168 | 0.0832 | 118.661 |
| R43 | CA | 0.9627 | 0.1932 | 56.599 |
| R43 | CO | 0.3004 | 0.0516 | 176.414 |

| | | | | |
|---|---|---|---|---|
| R43 | H | 0.3141 | 0.0473 | 7.991 |
| G44 | N | -0.732 | 0.1356 | 107.931 |
| G44 | CO | -0.7279 | 0.0887 | 173.213 |
| G44 | H | 0.0756 | 0.0235 | 8.178 |
| L45 | N | 0.3154 | 0.2641 | 121.959 |
| L45 | H | 0.9243 | 0.1137 | 8.361 |
| K46 | N | 0.7429 | 0.0402 | 123.48 |
| K46 | CA | 0.7697 | 0.5974 | 56.244 |
| K46 | CO | -1.5683 | 0.1172 | 174.582 |
| K46 | HA | -0.9243 | 0.324 | 4.304725 |
| V47 | N | -4.76 | 0.0648 | 122.427 |
| V47 | CA | -1.4717 | 0.2086 | 60.366 |
| V47 | CO | 1.6437 | 0.089 | 176.013 |
| V47 | H | -1.1571 | 0.1822 | 8.315 |
| C48 | N | -1.5898 | 0.0619 | 123.83 |
| C48 | CA | -2.0316 | 0.1631 | 54.325 |
| C48 | CO | 0.4475 | 0.091 | 173.52 |
| C48 | H | -0.2977 | 0.0333 | 8.772 |
| C48 | HA | -0.4336 | 0.0643 | 4.99944 |
| A49 | N | -1.6554 | 0.0809 | 125.577 |
| A49 | CA | 1.504 | 0.181 | 52.321 |
| A49 | CO | 1.3711 | 0.1132 | 176.226 |
| A49 | H | -1.1775 | 0.142 | 8.533 |
| D50 | N | 0.0047 | 2.4947 | 121.885 |
| D50 | H | -0.1974 | 0.012 | 8.441 |
| P51 | CO | 0.4459 | 0.0758 | 177.186 |
| Q52 | N | 5.3486 | 0.0469 | 118.002 |
| Q52 | CA | 1.2961 | 0.1332 | 56.466 |
| Q52 | CO | -1.022 | 0.0607 | 175.662 |
| Q52 | H | 0.155 | 0.0206 | 8.433 |
| A53 | N | -0.003 | 0.8999 | 124.115 |
| A53 | CO | -0.9638 | 0.1294 | 177.794 |
| A53 | H | 0.4901 | 0.0191 | 7.963 |
| T54 | N | -6.6514 | 0.0452 | 113.821 |
| T54 | H | -0.7506 | 0.0716 | 8.156 |
| W55 | N | 4.4177 | 0.077 | 122.321 |
| W55 | CO | -1.1239 | 0.115 | 176.196 |
| W55 | H | -0.135 | 0.0111 | 8.218 |

| | | | | |
|---|---|---|---|---|
| V56 | N | -3.0572 | 0.0468 | 121.369 |
| V56 | CO | -1.1368 | 0.1139 | 175.696 |
| V56 | H | 2.6243 | 0.3178 | 8.508 |
| R57 | N | 3.968 | 0.059 | 122.682 |
| R57 | CA | -2.5257 | 0.1148 | 56.787 |
| R57 | H | 0.4157 | 0.056 | 7.982 |
| D58 | N | 2.122 | 0.0525 | 120.936 |
| D58 | H | 0.0003 | 0.649 | 8.207 |
| V59 | CA | -3.7606 | 0.0307 | 62.968 |
| V59 | HA | 0.4031 | 0.0257 | 4.118123 |
| V60 | N | 2.2954 | 0.1797 | 122.998 |
| V60 | CO | -0.5545 | 0.0812 | 176.846 |
| R61 | N | 5.5122 | 0.0499 | 123.434 |
| R61 | CA | -2.6589 | 0.1024 | 56.702 |
| R61 | CO | -2.4068 | 0.0982 | 176.726 |
| R61 | H | 0.404 | 0.0312 | 8.3 |
| S62 | N | 2.0072 | 0.0484 | 116.542 |
| S62 | H | 0.2943 | 0.0143 | 8.093 |
| M63 | CA | -2.0217 | 0.1795 | 56.197 |
| D64 | N | -0.001 | 0.4951 | 121.555 |
| D64 | CA | -2.2031 | 0.1716 | 54.048 |
| D64 | CO | -1.6334 | 0.0971 | 175.971 |
| D64 | H | -0.2072 | 0.0191 | 8.395 |
| R65 | N | 1.5081 | 0.0544 | 121.806 |
| R65 | CA | -1.6576 | 0.1254 | 56.376 |
| R65 | CO | -1.2421 | 0.0784 | 176.263 |
| R65 | H | 0.2679 | 0.0107 | 8.241 |
| K66 | N | 2.0924 | 0.0556 | 122.117 |
| K66 | H | 0.3585 | 0.0175 | 8.374 |
| S67 | N | 1.2299 | 0.0513 | 116.587 |
| T69 | N | 0.0027 | 0.2171 | 114.177 |
| T69 | CA | 0.0003 | 0.4991 | 63.164 |
| R70 | N | 0.2029 | 0.1745 | 123.044 |
| R70 | CA | -0.0002 | 0.3147 | 56.456 |
| R70 | CO | -0.0065 | 2.9283 | 175.972 |
| N71 | N | 0.2721 | 0.101 | 119.766 |
| N71 | CO | 0.0021 | 40.8346 | 176.083 |
| N72 | N | -0.3333 | 0.0937 | 118.84 |

| | | | | |
|---|---|---|---|---|
| N72 | CA | 0.0011 | 1.3169 | 53.464 |
| N72 | CO | -0.0007 | 11.1529 | 174.977 |
| M73 | N | -0.0022 | 0.4251 | 120.234 |
| M73 | CA | -0.0009 | 0.913 | 55.69 |
| M73 | CO | 0.0723 | 0.3767 | 176.077 |
| I74 | N | 0.0046 | 0.4796 | 122.134 |
| I74 | CA | -0.0002 | 0.1756 | 61.121 |
| I74 | CO | 0.2517 | 0.1266 | 176.218 |
| Q75 | N | 0.3868 | 0.0528 | 125.291 |
| Q75 | CA | -0.003 | 2.199 | 55.575 |
| Q75 | CO | 0.1006 | 0.3238 | 175.901 |
| T76 | N | -0.3207 | 0.0569 | 116.51 |
| T76 | CA | 0 | 24.7953 | 61.905 |
| T76 | CO | 0.1595 | 0.1404 | 174.199 |
| K77 | N | 0.2232 | 0.0731 | 125.406 |
| P78 | CA | 0.0001 | 0.0414 | 63.072 |
| P78 | CO | 0.2148 | 0.0699 | 177.241 |
| T79 | N | 0.0043 | 0.3551 | 114.62 |
| T79 | CA | 0 | 24.0304 | 61.878 |
| T79 | CO | 0.3228 | 0.0564 | 175.416 |
| G80 | N | 0.1507 | 0.2417 | 111.329 |
| G80 | CO | 0.28 | 0.089 | 174.568 |
| T81 | N | 0.0021 | 0.2584 | 113.85 |
| T81 | CA | 0.0005 | 0.5842 | 61.89 |
| T81 | CO | 0.2625 | 0.0537 | 174.901 |
| Q82 | N | 0.2593 | 0.1211 | 123.289 |
| Q82 | CO | 0.1742 | 0.1182 | 175.943 |
| Q83 | N | 0.0058 | 0.9185 | 121.933 |
| Q83 | CO | 0.167 | 0.1845 | 176.089 |
| S84 | N | 0.2509 | 0.1136 | 117.698 |
| S84 | CA | -0.0019 | 1.5213 | 58.368 |
| S84 | CO | 0.0895 | 0.4024 | 174.802 |
| T85 | N | -0.3199 | 0.1031 | 115.367 |
| T85 | CA | 0 | 24.7855 | 61.636 |
| T85 | CO | 0.128 | 0.2854 | 174.303 |
| N86 | N | -0.4876 | 0.0725 | 120.844 |
| N86 | CO | 0.4267 | 0.0455 | 175.609 |
| T87 | N | -0.0416 | 0.5352 | 115.012 |

| | | | | |
|---|---|---|---|---|
| T87 | CA | 0 | 23.5994 | 61.936 |
| T87 | CO | 0.2822 | 0.0774 | 174.328 |
| A88 | N | 0.0106 | 0.7223 | 126.779 |
| A88 | CA | -0.0002 | 0.1554 | 52.596 |
| A88 | CO | 0.1293 | 0.1866 | 177.604 |
| V89 | N | 0.0013 | 0.2325 | 119.759 |
| V89 | CO | 0.0845 | 0.3633 | 176.219 |
| T90 | N | -0.0884 | 0.3204 | 118.906 |
| T90 | CA | 0.0003 | 0.2129 | 61.767 |
| T90 | CO | 0.1615 | 0.2541 | 174.336 |
| T92 | N | -0.0185 | 1.6586 | 114.284 |
| T92 | CA | 0.0003 | 0.193 | 61.524 |
| T92 | CO | 0.1704 | 0.111 | 174.151 |
| G93 | N | 0.001 | 0.1521 | 117.378 |

For residues with $\Delta\varpi$ values approaching zero, the fitting algorithm occasionally yielded large error estimates. However, visual inspection of these CEST profiles confirmed the absence of any minor exchange dips. Furthermore, the inclusion of these low $\Delta\varpi$ values did not affect the calculation of the ES structure.

**Table S2:** List of NMR samples used in this work.

| Sample no | Variant | Labelling | Concentration (mM) | Experiments | Remarks |
|---|---|---|---|---|---|
| 1 | WT | $^{15}N$ | 1.1 | $^{15}N$ CEST, $^{15}N$ CPMG | To obtain excited state chemical shifts and $p_E$,$k_{EX}$ |
| 2 | WT | $^{15}N$ | 0.5 | $^{15}N$ CEST | Concentration dependent CEST |
| 3 | WT | $^{13}C$,$^{15}N$ | 1.9 | $^{13}C\alpha$ CEST, $^{13}C$\` CEST, $^{13}C$\` CPMG, $^{1}HN$ CEST, $^{1}HN$ CPMG, $^{15}N$ CEST | To obtain multinuclear backbone chemical shifts for the excited state, concentration dependent CEST |
| 4 | WT | $^{13}C$,$^{15}N$,$^{2}H$ | 0.99 | $^{13}C\alpha$ CEST (HA-CA based) | Sample in 100% $D_2O$ buffer. To obtain $^{13}C\alpha$ chemical shifts for the excited state |
| 5 | WT | $^{13}C$,$^{15}N$,$^{2}H$ | 0.82 | $^{1}H\alpha$ CPMG, $^{1}H\alpha$ $R_{1\rho}$ | To obtain the magnitude of $^{1}H\alpha$ chemical shifts along with their signs |
| 6 | R61A | $^{15}N$ | 0.9 | $^{15}N$ CEST, $^{15}N$ CPMG | To obtain $^{15}N$ chemical shifts and $p_E$,$k_{EX}$ |
| 7 | V59A | $^{15}N$ | 1.3 | $^{15}N$ CEST, $^{15}N$ CPMG | To obtain $^{15}N$ chemical shifts and $p_E$,$k_{EX}$ |
| 8 | S62Y | $^{15}N$ | 1.2 | $^{15}N$ CEST, $^{15}N$ CPMG | To obtain $^{15}N$ chemical shifts and $p_E$,$k_{EX}$ |
| 9 | S13P | $^{15}N$ | 0.99 | $^{15}N$ CEST, $^{15}N$ CPMG | To obtain $^{15}N$ chemical shifts and $p_E$,$k_{EX}$ |
| 10 | CC3 | $^{15}N$ | 1.3 | $^{15}N$ CEST, $^{15}N$ CPMG | To obtain $^{15}N$ chemical shifts and $p_E$,$k_{EX}$ |

| | | | | | |
|---|---|---|---|---|---|
| 11 | Δ(69-93) | $^{15}N$ | 1.2 | $^{15}N$ CEST, $^{15}N$ CPMG | To obtain $^{15}N$ chemical shifts and $p_E$,$k_{EX}$ |
| 12 | V59P | $^{15}N$ | 0.45 | $^{1}H$-$^{15}N$ HSQC | - |
| 13 | V56G | $^{15}N$ | 0.32 | $^{1}H$-$^{15}N$ HSQC | - |
| 14 | Δ(55-93) | $^{15}N$ | 0.032 | $^{1}H$-$^{15}N$ HSQC | - |
| 15 | CC5/V12R | $^{15}N$ | 0.14 | $^{1}H$-$^{15}N$ HSQC | - |

**Table S3:** Experimental parameters for CEST experiments on WT and mutant Ltn. Sample IDs are taken from Table S2.

| Sample No. | Experiment | B1 (Hz) | $T_{ex}$ (s) | Sweep (Hz) | No. of planes | Offset spacing (Hz) |
|---|---|---|---|---|---|---|
| 1 | $^{15}$N CEST | 15.3 | 0.4 | -810 to 810 | 110 | 15 |
| | | 30.4 | 0.4 | -810 to 810 | 56 | 30 |
| | | 48.5 | 0.4 | -810 to 830 | 43 | 40 |
| 2 | $^{15}$N CEST | 30.2 | 0.4 | -810 to 810 | 56 | 30 |
| | | 48.2 | 0.4 | -810 to 830 | 43 | 40 |
| 3 | $^{15}$N CEST | 30.3 | 0.4 | -810 to 810 | 56 | 30 |
| | $^{13}$Cα CEST | 33.0 | 0.3 | -1250 to 1480 | 93 | 30 |
| | $^{13}$CO CEST | 31.0 | 0.3 | -900 to 900 | 62 | 30 |
| | $^{1}$HN CEST | 25 | 0.5 | 690 to 4230 | 120 | 30 |
| 4 | $^{13}$Cα CEST | 33 | 0.25 | -2400 to 2000 | 112 | 40 |
| | | 43.8 | 0.25 | -2350 to 1950 | 88 | 50 |
| | | 54.6 | 0.25 | -2400 to 1980 | 75 | 60 |
| 6 | $^{15}$N CEST | 26.6 | 0.4 | -960 to 960 | 66 | 30 |
| | | 42.1 | 0.4 | -1120 to 1120 | 58 | 40 |
| 7 | $^{15}$N CEST | 21.2 | 0.4 | -960 to 965 | 79 | 25 |
| | | 31.6 | 0.4 | -1035 to 1030 | 61 | 35 |
| 8 | $^{15}$N CEST | 26.1 | 0.4 | -990 to 990 | 68 | 30 |
| | | 42.1 | 0.4 | -1000 to 1000 | 52 | 40 |
| 9 | $^{15}$N CEST | 26.7 | 0.4 | 930 to 930 | 64 | 30 |
| 10 | $^{15}$N CEST | 26.0 | 0.4 | -1200 to 1020 | 76 | 30 |
| | | 41.6 | 0.4 | -980 to 980 | 51 | 40 |
| 11 | $^{15}$N CEST | 31.4 | 0.35 | -700 to 595 | 39 | 35 |
| | | 50.5 | 0.35 | -755 to 620 | 27 | 55 |

**Table S4:** Experimental parameters for CPMG experiments on WT and mutant Ltn. Sample IDs are taken from Table S2.

| Sample No. | Experiment | $B_0$ (MHz) | CPMG $T_{ex}$ (ms) | Highest CPMG frequency (Hz) | No. of planes |
|---|---|---|---|---|---|
| 1 | $^{15}$N CPMG | 700 | 30 | 1000 | 21 |
| | $^{15}$N CPMG (10 °C) | 700 | 30 | 1000 | 21 |
| 3 | $^{13}$C\` CPMG | 700 | 40 | 975 | 22 |
| | $^{1}$HN CPMG | 700 | 20 | 2000 | 25 |
| | | 600 | 20 | 2000 | 24 |
| 5 | $^{1}$Hα CPMG | 800 | 18 | 1888 | 19 |
| | | 600 | 18 | 1888 | 19 |
| 6 | $^{15}$N CPMG | 700 | 30 | 1000 | 21 |
| 7 | $^{15}$N CPMG | 700 | 30 | 1000 | 21 |
| 8 | $^{15}$N CPMG | 700 | 30 | 1000 | 21 |
| 9 | $^{15}$N CPMG | 800 | 30 | 1000 | 22 |
| 10 | $^{15}$N CPMG | 700 | 30 | 1000 | 21 |
| 11 | $^{15}$N CPMG | 600 | 30 | 1000 | 19 |

**Table S5:** List of experimental parameters used in $^{1}H\alpha$ $R_{1\rho}$ experiments.

| Residue | $\lvert\delta_G\rvert$ (Hz) | $\nu_1$ (Hz) |
|---|---|---|
| V21 | 185 | 90 |
| Y27 | 625 | 185 |
| F39 | 285 | 115 |
| I40 | 145 | 70 |
| K42 | 100 | 65 |
| K46 | 530 | 163 |
| C48 | 185 | 95 |
| V59 | 205 | 105 |
